# Calcium buffering tunes intrinsic excitability of spinal dorsal horn parvalbumin-expressing interneurons: A computational model

**DOI:** 10.1101/2023.03.05.531043

**Authors:** Xinyue Ma, Loïs Miraucourt, Haoyi Qiu, Reza Sharif-Naeini, Anmar Khadra

**Author notes:** Co-corresponding authors. Corresponding author email address: Reza Sharif-Naeini, Anmar Khadra.

## Abstract

Parvalbumin-expressing interneurons (PVINs) play a crucial role within the dorsal horn of the spinal cord by preventing touch inputs from activating pain circuits. After nerve injury, their output is decreased via mechanisms that are not fully understood. In this study, we show that PVINs from nerve-injured mice change their firing pattern from tonic to adaptive. To examine the ionic mechanisms responsible for this decreased output, we employed a reparametrized Hodgkin-Huxley (HH) type model of PVINs, which predicted (1) the firing pattern transition is due to an increased contribution of small conductance calcium-activated potassium (SK) channels, enabled by (2) impairment in intracellular calcium buffering systems. Analyzing the dynamics of the HH-type model further demonstrated that a generalized Hopf bifurcation differentiates the two types of state transitions observed in the transient firing of PVINs. Importantly, this predicted mechanism holds true when we embed the PVINs model within the neuronal circuit model of the spinal dorsal horn. To experimentally validate this hypothesized mechanism, we used pharmacological modulators of SK channels and demonstrated that (1) tonic firing PVINs from naïve mice become adaptive when exposed to an SK channel activator, and (2) adapting PVINs from nerve-injured mice return to tonic firing upon SK channel blockade. Our work provides important insights into the cellular mechanism underlying the decreased output of PVINs in the spinal dorsal horn after nerve injury and highlights potential pharmacological targets for new and effective treatment approaches to neuropathic pain.

**Significant Statement:** Parvalbumin-expressing interneurons (PVINs) exert crucial inhibitory control over A *β* fiber-mediated nociceptive pathways at the spinal dorsal horn. The loss of their inhibitory tone leads to neuropathic symptoms, like mechanical allodynia, via mechanisms that are not fully understood. This study identifies the reduced intrinsic excitability of PVINs as a potential cause for their decreased inhibitory output in nerve-injured condition. Combining computational and experimental approaches, we predict a calcium-dependent mechanism that modulates PVINs’ electrical activity following nerve injury: a depletion of cytosolic calcium buffer allows for the rapid accumulation of intracellular calcium through the active membranes, which in turn potentiates SK channels and impedes spike generation. Our results therefore pinpoint SK channels as interesting therapeutic targets for treating neuropathic symptoms.

## Introduction

Neuropathic pain is a chronic debilitating disease that can be induced by nerve injury or occur spontaneously in the somatosensory system. It affects ∼10% of the population (Van Hecke et al., 2014) with symptoms including mechanical allodynia (Baron, 2009), a painful response to innocuous mechanical stimuli. This symptom severely reduces the life quality of patients (Anon, 2019), the care of which has become a significant economic burden to society (Guerriere et al., 2010).

Neuropathic pain has been linked to an imbalance of inhibitory and excitatory signaling in dorsal horn of the spinal cord (Gwak and Hulsebosch, 2011), where the sensory information is relayed to the central nervous system. At this site, the immunoreactivity for the inhibitory neurotransmitter gamma-aminobutyric acid (GABA) was shown to be decreased after nerve injury (Ibuki et al., 1996; Eaton et al., 1998) and reversely, pharmacological blockade of inhibitory synaptic transmission in the naïve animals can induce mechanical allodynia (Yaksh, 1989). Importantly, the reduction in inhibitory tone after nerve injury was not associated with a loss of inhibitory neurons (Bennett and Xie, 1988; Petitjean et al., 2015). Other possibilities, including reduced electrical activity of inhibitory interneurons (Boyle et al., 2019; Qiu et al., 2022) or a decrease in their synaptic connections (Petitjean et al., 2015) may be involved.

A pivotal element within the spinal microcircuits in the dorsal horn is a subset of inhibitory parvalbumin-expressing interneurons (PVINs). PVINs function as gate-keepers in preventing touch information from activating pain-specific projection neurons (Petitjean et al., 2015). PVINs have multiple post-synaptic targets in the dorsal horn, one of which is the protein kinase Cγ (PKCγ)-containing excitatory interneurons located in the superficial layer of the dorsal horn (Petitjean et al., 2015; Boyle et al., 2019). The PKCγ interneurons are involved in the polysynaptic transmission of low-intensity touch inputs to projection neurons and are normally under the inhibitory control of PVINs (Miraucourt et al., 2007). Boyle et. al. previously showed that the alteration of PVINs’ excitability can impact their inhibitory output (Boyle et al., 2019), but the ionic mechanism underlying the changes of their intrinsic electrical properties remains incompletely understood.

In this study, we demonstrated that after nerve injury, PVINs change their firing pattern from tonic to adaptive. To explore the mechanisms underlying the change in PVIN firing properties, we combined computational and experimental approaches. Our results point to a calcium-dependent mechanism that is critical to PVIN’s electrical activity and may be impaired after nerve injury. This finding was obtained by initially reparametrizing a previously developed conductance-based Hodgkin-Huxley (HH) type model (Bischop et al., 2012; Orduz et al., 2013) to reproduce the firing properties of PVINs from naïve mice, followed by identifying the role of small conductance calcium-activated potassium (SK) channels in inducing the transition in the firing pattern of PVINs after nerve injury. This prediction was further confirmed experimentally using pharmacological agents that activate or block SK channels. In brief, this mechanism suggests that, in fast-spiking PVINs, calcium buffer depletion allows for the rapid accumulation of intracellular calcium through their depolarized membranes, which in turn potentiates SK channels and decreases their spiking behavior.

## Materials and methods

### Animals

All mice were kept under pathogen-free conditions, and cages were maintained in ventilated racks in temperature (20–21 ℃) and humidity (55%) controlled rooms on a 12 h light/dark cycle, with food and water provided *ad libitum*. All experimental procedures were approved by the Animal Care and Use Committee at McGill University, in accordance with the regulations of the Canadian Council on Animal Care. The PV^cre^;tdTom mice were generated by crossing commercially available PV^cre^ mice (JAX, stock #017320, https://www.jax.org/strain/017320) with Ai14 tdTomato reporter (JAX, stock#007914, https://www.jax.org/strain/007914).

### Chronic Constriction Injury Surgery

Unilateral sciatic nerve chronic constriction injury (CCI) was done as previously described (Austin et al., 2012). Three ligatures (6-0 Perma-Hand silk suture; Ethicon) were placed around the sciatic nerve bundle and loosely closed to construct but not arrest epineural blood flow. Mice developed robust mechanical allodynia that becomes chronic after two weeks and can persist for up to 8 weeks post-surgery. To compromise the variation of the surgery, the CCI surgery was all performed by one experienced experimenter.

### Slice Preparation

Adult male mice (12-16 weeks old) were anesthetized with 5% isofluorane and perfused transcardially with ice-cold oxygenated (95% O2, 5% CO2) N-Methyl-D-glutamine-based artificial cerebrospinal fluid (NMDG-ACSF) solution (93 NMDG, 93 HCl 12M, 2.5 KCl, 1.25 NaH2PO4, 30 NaHCO3, 20 HEPES, 25 glucose, 2 thiourea, 5 Na-L-ascorbate, 3 Na-pyruvate, 10 N-acetyl-L cysteine, 0.5 CaCl2/2H2O, and 10 MgSO4/7H2O, in mM; 300±10 mOsm; pH 7.3–7.4). The lumbar spinal segment was rapidly removed and immersed in ice-cold NMDG-ACSF. The ventral roots were removed and 300 µm transverse slices were cut with a vibratome (Leica VT1000S). Slices were transferred to a submerged chamber containing HEPES-based recovery ACSF (92 NaCl, 2.5 KCl, 1.25 NaH2PO4, 30 NaHCO3, 20 HEPES, 25 Glucose, 2 Thiourea, 5 Na-ascorbate, 3 Na- pyruvate, 2 CaCl2, 2 MgSO4, in mM; 300±10 mOsm, pH 7.4) for at least 10 minutes at 36 ℃, equilibrated with 95% O2 and 5% CO2. At the end of the slicing period, the HEPES-based recovery ACSF wet let to equilibrate at room temperature for at least one hour.

### Electrophysiology

Slices were transferred individually to a recording chamber and continuously superfused with oxygenated ACSF (126 NaCl, 26 NaHCO3, 2.5 KCl, 1.25 NaH2PO4, 2 CaCl2, 2 MgCl2, and 10 glucose, in mM; bubbled with 95% O2 and 5% CO2; pH 7.3; osmolarity, 300±10 mOsm measured, room temperature, 2 ml/min). Patch pipettes were pulled from borosilicate glass capillaries (Harvard Apparatus) with a P-97 puller (Sutter Instruments). They were filled with a solution containing 135 K-gluconate, 6 NaCl, 0.1 EGTA, 10 HEPES, 2 MgCl2, 2 Mg-ATP, and 0.8 Na-GTP (in mM; pH 7.4, adjusted with KOH; osmolarity, 300±10 mOsm, adjusted with sucrose) and had final tip resistances of 6–8 MΩ for whole cell recording. Holding potentials were applied to correct a liquid junction potential of −15.9 mV. Neurons were viewed by an upright microscope (Olympus) with a 40X water-immersion objective, infrared differential interference contrast (IR-DIC) and fluorescence. Recordings were made from identified PV-neurons expressing the tdTomato. The whole-cell patch configuration can potentially affect the endogenous calcium and calcium buffers and this artifact was minimized by recording right after we break through the cell. Data was acquired with pClamp 10 suite software (Molecular Devices) using MultiClamp 700B patch-clamp amplifier and Digidata 1440A (Molecular Devices). Recordings were low-pass filtered on-line at 1 kHz, digitized at 20 kHz. For SK current manipulation, 200 nM apamin or 0.3 mM 1-EBIO was added in the bath solution.

We applied 3 standardized current-clamp protocols per cell recording. A gap-free 90-s long recording with no injection current (I=0) was used to measure the resting membrane potential (mV). A 1-sec long step current injection from -200 pA to 200 pA with a step size of 50 pA, and a 1-sec long ramp current injection from 0 pA to 400 pA with a step size of 50 pA were used to measure the active membrane properties including input resistance (MΩ), rheobase (pA), spike frequency (Hz), spike threshold (mV), half-width (ms), action potential peak (mV) and trough (mV), half-width (ms), action potential rise and fall time (ms), and after-hyperpolarization time (ms).

### Data analysis

All the passive and active membrane properties were quantified as commonly defined. Resting membrane potential was measured as the average of voltage in the last 30-s period from a 90-s recording with no injection current. Input resistance was obtained from the slope of the I-V curve in response to hyperpolarized injection currents. Whole-cell capacitance was determined by the amplifier’s automated whole-cell compensation function. Rheobase was defined as the minimal injected step current value needed to evoke a single action potential. We plotted the latency of the first spike against the injected step current value, fit the datapoint to power function *I*_*app*_ = *t*^−*a*^ + *b* where the rheobase was measured as the bottom asymptote y=b that cross Y-axis. The spike threshold, measured as the point at which 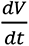 deviated from the mean of the baseline by 3 standard deviations. Here, the baseline is defined as the 5 ms long voltage response before 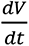 reaches 3 mV/ms, which was chosen empirically so that the detected spike threshold lies at the elbow of spike depolarization phase of the AP cycle. AP peak is the maximum potential and AP trough is the minimum potential in the interval between two consecutive AP peaks. Half-width is the AP duration at the membrane voltage halfway between the spike threshold and AP peak. AP rise and fall times were calculated between 10% and 90% of the voltages from spike threshold to AP peak. The area of medium after-hyperpolarization potential (mAHP) was measured as the area between the resting voltage and the voltage within 200 ms after the test stimulus. The amplitude of mAHP is the difference between the resting voltage and the minimal voltage within 200 ms after the test stimulus. Firing frequencies were measured as the reciprocal of the inter-spike intervals. Its time course was well fitted by a single exponential function 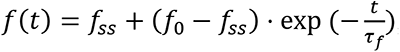, where *τ_f_* denotes the spike frequency decay rate. A transiently firing CCI recording was identified as an “on-off switching” if any spike peak in that recording decreases ≥20 mV compared to its formal spike; otherwise, it was identified as “damped oscillation”. The voltage sag ratio is measured as the voltage sag amplitude deflection divided by the steady-state voltage deflection, to normalize for input resistance differences between cells, where “deflection” refers to the voltage difference from the baseline voltage before the hyperpolarizing current. All the voltage properties were not adjusted with the liquid junction potential of -15.9 mV.

### PVIN Neuron Computational Model

A modified HH type model was adopted from Bischop et. al. 2012 (Bischop et al., 2012). The model from Bischop et. al. 2012 was reparametrized to accurately capture the cycles of action potentials in (*V*, *V*)-plane and firing frequencies of the naïve PVINs recordings (**Extended Fig. 2-1 and 2-2**). The model presented in this work is available for public download in ModelDB (http://modeldb.yale.edu/267614, read-only password: 584953937798738).

The resulting 5-dimensional HH type model for PVIN is given by

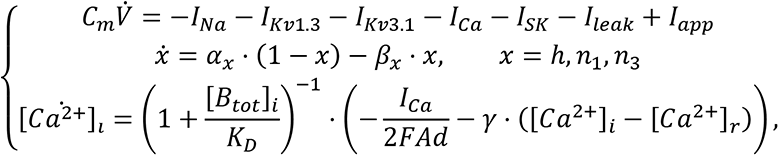

where *C*_*m*_ and *V* are the membrane capacitance and potential of the PVIN, [*Ca*^2+^]_*i*_ and [*B*_*tot*_]_*i*_ are the intracellular concentration of calcium and calcium buffers. According to Eq. (1), the model consists of six ionic currents, including

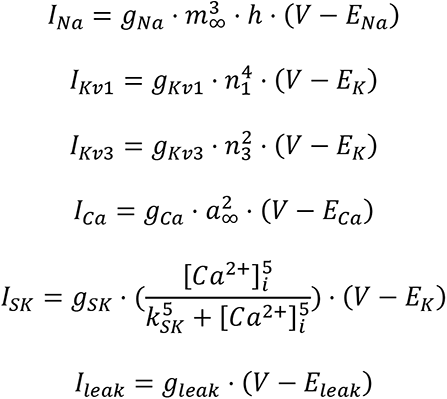

where *g*_*x*_ and *E*_*x*_ (*x* = *Na*, *K*, *Ca*, *Leak*) are the maximal ionic conductance and the Nernst potential of each current, respectively, *n*_1_ and *n*_3_ are the activation variables of *I*_*Kv*1_ and *I*_*Kv*3_, respectively, and *m*_∞_ and *a*_∞_ are the steady state activations of *I*_*Na*_ and *I*_*Ca*_, respectively. The kinetics of the various ionic currents obtained from fitting the AP cycles to data are described below with figures comparing the voltage-dependent steady state activation/inactivation functions and time constants before (grey) and after (black) parameter tuning. The full list of model parameters is provided in **Table 1**.

**Table 1.** PVIN model parameters.

| Parameter | Value | Unit | Definition | Reference |
| --- | --- | --- | --- | --- |
| $C_m$ | 30 | pF | Membrane capacitance | (Bischof et al., 2012; Orduz et al., 2013) |
| $g_{Na}$ | 300 | nS | Maximum $Na^+$ conductance | Estimated |
| $g_{Kv1}$ | 15 | nS | Maximum Kv1 $K^+$ conductance | Estimated |
| $g_{Kv3}$ | 180 | nS | Maximum Kv3 $K^+$ conductance | Estimated |
| $g_{SK}$ | 10 | nS | Maximum SK $K^+$ conductance | Estimated |
| $g_{Ca}$ | 8 | nS | Maximum $Ca^{2+}$ conductance | Estimated |
| $g_{leak}$ | 8 | nS | Maximum leak conductance | Estimated |
| $E_{Na}$ | 58 | mV | $Na^+$ Nernst potential | Estimated |
| $E_K$ | -80 | mV | $K^+$ Nernst potential | Estimated |
| $E_{Ca}$ | 68 | mV | $Ca^{2+}$ Nernst potential | Estimated |
| $E_{leak}$ | -50 | mV | Leak Nernst potential | Estimated |
| $\gamma$ | 0.01 | $ms^{-1}$ | $Ca^{2+}$ recovery rate | Estimated |
| $k_{SK}$ | 0.8 | $\mu M$ | Calcium sensitivity of SK channels | Estimated |
| A | 3000 | $\mu\text{m}^2$ | Cell surface area | (Bischof et al., 2012; Orduz et al., 2013) |
| d | 0.1 | $\mu\text{m}$ | Shell thickness | (Bischof et al., 2012; Orduz et al., 2013) |
| $[\text{Ca}^{2+}]_r$ | 0.07 | $\mu\text{M}$ | Resting cytosolic $\text{Ca}^{2+}$ concentration | (Bischof et al., 2012; Orduz et al., 2013) |
| $K_D$ | 0.1 | $\mu\text{M}$ | Buffer affinity for $\text{Ca}^{2+}$ | (Schwaller et al., 2002) |

1. Fast transient Na^+^ current (*I*_*Na*_):

The gating of *I*_*Na*_ is governed by both the steady state activation function

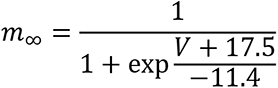

and the inactivation variable ℎ satisfying the dynamic equation

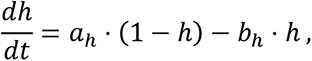

where

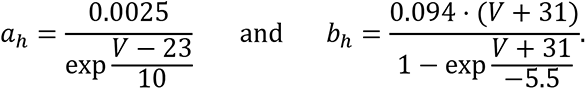

2. Slow delayed-rectifier K^+^ current (*I*_*Kv*1.3_)

The gating of *I*_*Kv*1.3_ is governed by the activation variable *n*_1_, satisfying the dynamic equation

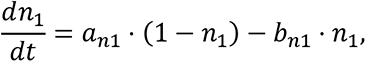

where

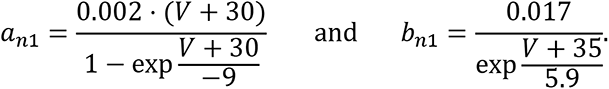

3. Fast delayed-rectifier K^+^ current (*I*_*Kv*3.1_)

The gating of *I*_*Kv*3.1_ is governed by the activation variable *n*_3_, satisfying the dynamic equation

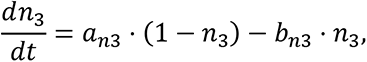

where

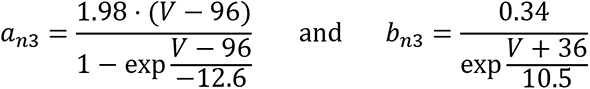

4. High-voltage activated Ca^2+^ current (*I*_*Ca*_)

The gating of *I*_*Ca*_ is governed by the steady state activation function

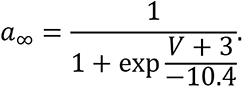

The model was stimulated by the same step current protocol for data acquisition. To study the dynamics of PVIN under realistic conditions, we stimulated PVIN models with a 3-sec Poisson-distributed presynaptic current mimicking the presynaptic inputs from the primary afferents Aβ fibers. The frequency of the presynaptic input corresponds to the firing rate of *Aβ* fibers in response to varied mechanical stimulation as previously reported (Walcher et al., 2018).

### PKC*γ* Neuron Computational Model

We adopted a HH type model for the somatic compartment of excitatory PKC*γ*IN at the spinal dorsal horn using the formalism from Medlock et. al. 2022. The model is given by

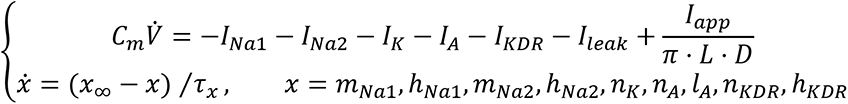

The ionic currents of the model include HH-type 1 *Na*^+^ current (*I*_*Na*1_, Traub and Miles, 1991), HH-type 2 *Na*^+^ current (*I*_*Na*2_, Hodgkin and Huxley, 1952), HH-type *K*^+^ current (*I*_*K*_, Hodgkin and Huxley, 1952), delayed-rectifier *K*^+^ current (*I*_*KDR*_, Safronov et al. 2000), A-type *K*^+^ current (*I*_*A*_, Whyment et al. 2011) and a non-specific leak current (*I*_*leak*_):

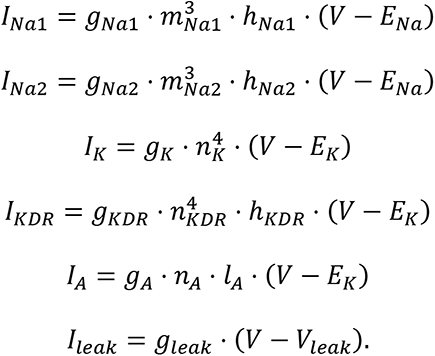

The various parameters introduced in these currents are defined are before in the PVIN model. The kinetics of the varied ionic currents are supplied in the freely available code in ModelDB (http://modeldb.yale.edu/267056). Parameters are summarized in **Table 2**.

**Table 2.** PKC*γ* model parameters (Medlock et al., 2022).

| Parameter | Value | Unit | Definition |
| --- | --- | --- | --- |
| $C_m$ | 1 | $\mu\text{F}/\text{cm}^2$ | Membrane capacitance |
| $g_{\text{Na1}}$ | 0.1625 | $\text{mS}/\text{cm}^2$ | Maximum HH-type 1 $\text{Na}^+$ conductance |
| $g_{\text{Na2}}$ | 85.48 | $\text{mS}/\text{cm}^2$ | Maximum HH-type 2 $\text{Na}^+$ conductance |
| $g_K$ | 4.3 | $\text{mS}/\text{cm}^2$ | Maximum HH-type $\text{K}^+$ conductance |
| $g_A$ | 10.9 | $\text{mS}/\text{cm}^2$ | Maximum A-type $\text{K}^+$ conductance |
| $g_{\text{KDR}}$ | 3.111 | $\text{mS}/\text{cm}^2$ | Maximum delayed-rectifier $\text{K}^+$ conductance |
| $g_{\text{leak}}$ | 0.96 | $\text{mS}/\text{cm}^2$ | Maximum leak conductance |
| $E_{\text{Na}}$ | 50 | mV | $\text{Na}^+$ Nernst potential |
| $E_K$ | -70 | mV | $\text{K}^+$ Nernst potential |
| $E_{\text{leak}}$ | -65 | mV | Leak Nernst potential |
| L | -60 | $\mu\text{m}$ | Soma length |
| D | 66 | $\mu\text{m}$ | Soma diameter |
| T | 23 | $^{\circ}\text{C}$ | Temperature |

### Synaptic Connectivity

Synaptic connectivity of the touch sensory neural circuit at spinal dorsal horn was based on recent experimental data (Petitjean et al., 2015). A*β* fibers excite both PVIN and PKC*γ* interneuron mediated by AMPA and NMDA receptors, while PKC *γ* interneuron also receives inhibitory synaptic inputs from PVIN by glycine and *GABA*_*A*_ receptors.

A synaptic current *I*_*syn*_ was added to Eqs. (1) and (3) describing the voltage dynamics of PVIN and PKC*γ* neurons. This synaptic current is given by

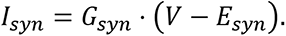

The synapse maximum conductance *G*_*syn*_ was modeled using two exponential functions with a rise (*τ*_*syn*,*r*_) and decay (*τ*_*syn*,*d*_) time constant, a synaptic strength *A*_*syn*_ defining the amplitude, and a factor *f*_*syn*_ to normalize the peak at 1. In other words,

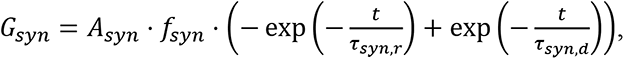

where *τ*_*syn*,*r*_ < *τ*_*syn*,*d*_, and *syn* ∈ {*AMDA*, *NMDA*, *glycine*, *GABA*_*A*_}. The rise and decay time constants (in milliseconds) of AMPA, NMDA, glycine and *GABA*_*A*_ are 0.1/5, 2/100, 0.1/10, 0.1/20, respectively (Medlock, et. al. 2022). The excitatory Nernst potential *E_exc_* is 0 mV and the inhibitory Nernst potential *E*_*int*_ is -70 mV. The magnesium block of NMDA receptor *M*_*g*_*block*__(*V*) is given by

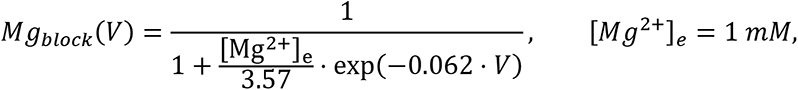

and is multiplied by the maximal conductance *G*_*syn*_. The strengths of synaptic connections were tuned to generate realistic electrophysiological responses to stimulation of primary afferent fibers and are given in **Table 3**.

**Table 3.** The strength parameters of synaptic connections.

| Connection | Receptor | Synaptic Strength $A_{\text{syn}}$ (nS) |
| --- | --- | --- |
| $(A\beta \rightarrow PKC\gamma IN)_e$ | AMPA | 5.5 |
|  | NMDA | 2 |
| $(A\beta \rightarrow PVIN)_e$ | AMPA | 2 |
|  | NMDA | 3 |
| $(PVIN \rightarrow PKC\gamma IN)_i$ | Glycine | 5.32 |
|  | GABA<sub>A</sub> | 10 |

The *Aβ* fiber-driven synaptic input was generated as 3-sec-long Poisson distributed spikes, the frequencies of which depends on the responses of the primary afferents *Aβ* -fibers to different mechanical (Walcher et al., 2018; Medlock et al., 2022). Accordingly, the firing rates of *Aβ* fibers were scaled linearly between 2-20 Hz for low mechanical forces ranging between 5-50 mN and plateauing at forces ≥ 50 mN. We therefore use 5 Hz and 20 Hz to imitate the firing rates of *Aβ* fibers in response to innocuous and noxious mechanical stimuli.

### Software and Numerical Simulations

Numerical simulations were performed using a code written in MATLAB R2022a (MathWorks). The one-parameter bifurcation diagrams were computed using XPPAUT (a freeware by Bard Ermentrout available online: http://www.math.pitt.edu/~bard/xpp/xpp.html), whereas the two-parameter bifurcation diagrams were generated by the MATCONT package.

### Code accessibility

All codes for the modeling are available for public download in ModelDB (http://modeldb.yale.edu/267614, read-only password: 584953937798738).

### Statistical Analysis

Data were expressed as mean±SEM, with n being the number of cells. To evaluate the difference between two groups, the normal-distributed data was compared using a t test, whereas a Mann- Whitney U test was used when data was not normally distributed. For multiple groups, ANOVA was used. P≤0.05 (*) was considered to indicate a statistically significant difference, with p<0.01 (**), p<0.001 (***) and p<0.0001 (****) also be noted. Statistical analysis was performed in GraphPad Prism 8.0.2.

## Results

### Nerve injury induces changes in the firing pattern and active membrane properties of PVINs

PVINs in the spinal dorsal horn exert crucial inhibitory control to the A*β*-mediated transduction of touch-sensory information. A loss of inhibitory tone from PVINs has been associated with nerve injury-induced mechanical allodynia. To characterize the impact of nerve injury on PVINs membrane properties, we performed targeted whole-cell patch-clamp recordings in acute spinal cord slice from PV^cre^;tdTom mice. Current-clamp recordings were performed in laminae IIinner and III tdTomato-positive interneurons in naïve (“naïve-PVINs”) and nerve injured (chronic constriction injury; “CCI-PVINs”) mice. The passive and active membrane properties were summarized in **Table 4**.

**Table 4.** Electrophysiological features of PVINs in the naïve (n=24) and CCI (n=30) mice. Data are mean±SEM. Unpaired two-tailed t test, *p<0.1, **p<0.01, ***p<0.001, ****p<0.0001.

| Properties |  | naïve | CCI | p value |
| --- | --- | --- | --- | --- |
| Resting membrane potential (mV) | | -52.79±1.26 | -49.30±1.69 | 0.1202 |
| Input resistance (MΩ) | | 274.09±15.71 | 241.22±23.19 | 0.2540 |
| Whole cell capacitance (pF) | | 18.17±3.16 | 12.66±1.35 | 0.1011 |
| Rheobase (pA) | | 27.72±4.00 | 28.94±3.33 | 0.8144 |
| Voltage sag ratio | | 1.43±0.04 | 1.50±0.06 | 0.3038 |
| AP peak (mV) | 1st spike | 18.89±1.61 | 22.01±1.54 | 0.1702 |
| | 3rd spike | 14.92±1.19 | 9.42±1.58 | 0.0097** |
|  | 5th spike | 13.96±1.23 | 4.80±1.92 | 0.0003*** |
| AP trough (mV) | 1st spike | -42.96±1.97 | -45.89±1.66 | 0.2579 |
|  | 3rd spike | -39.35±2.01 | -39.49±1.84 | 0.9611 |
|  | 5th spike | -38.52±2.01 | -37.36±1.98 | 0.6840 |
| Spike threshold (mV) | 1st spike | -34.40±1.10 | -37.95±1.61 | 0.0871 |
|  | 3rd spike | -28.48±1.28 | -29.21±1.35 | 0.7014 |
|  | 5th spike | -27.19±1.27 | -26.61±1.67 | 0.7903 |
| Half-width (ms) | 1st spike | 0.96±0.06 | 1.22±0.08 | 0.0145* |
|  | 3rd spike | 1.17±0.08 | 1.87±0.15 | 0.0004*** |
|  | 5th spike | 1.25±0.09 | 2.29±0.20 | 3e-5**** |
| AP rise time (ms) | 1st spike | 0.46±0.02 | 0.64±0.05 | 0.0015** |
|  | 3rd spike | 0.66±0.04 | 1.06±0.07 | 9e-6**** |
|  | 5th spike | 0.73±0.04 | 1.38±0.10 | 1e-6**** |
| AP fall time (ms) | 1st spike | 0.90±0.07 | 1.21±0.12 | 0.0377* |
|  | 3rd spike | 0.97±0.08 | 1.49±0.15 | 0.0058** |
|  | 5th spike | 1.01±0.08 | 1.62±0.16 | 0.0014** |
| AHP time (ms) | 1st spike | 1.02±0.06 | 1.30±0.12 | 0.0474* |
|  | 3rd spike | 1.38±0.07 | 1.75±0.14 | 0.0250* |
|  | 5th spike | 1.60±0.08 | 3.02±0.50 | 0.0132* |

In other nerve injury models where denervation was made, PVINs in denervated regions of the spinal dorsal horn were reported to be less active, producing less spikes in response to step currents (Boyle et al., 2019). However, a common factor critical to neuronal information coding is the firing frequency (Ainsworth et al., 2012). This is particularly important for inhibitory neurons (Ferrante et al., 2009; Bastos et al., 2012) such as PVINs, whose output prevents Aβ fibers from activating pain pathways (Medlock et al., 2022). We fitted spike frequencies to a single exponential decay function to the data, where the time constant denotes the adaptation rate in frequency (**Fig. 1A-B**). The naïve-PVINs exhibited predominantly tonic firing during a 1-s stable current stimulus (18 out 24 cells) with a slow frequency adaptation at a rate of 1.31±0.21 s (n=24). This is consistent with the characteristics of PVINs as fast-spiking interneurons with little spike frequency adaptation. Interestingly, nerve injury caused PVINs to exhibit a more transient firing pattern. In this context, PVINs from CCI mice failed to maintain tonic firing (**Fig. 1D**), with their frequencies decreasing steeply at a rate of 0.42 ± 0.14 s (n=30, **Fig. 1C**), demonstrating fast spike frequency adaptation.

**Figure 1.**
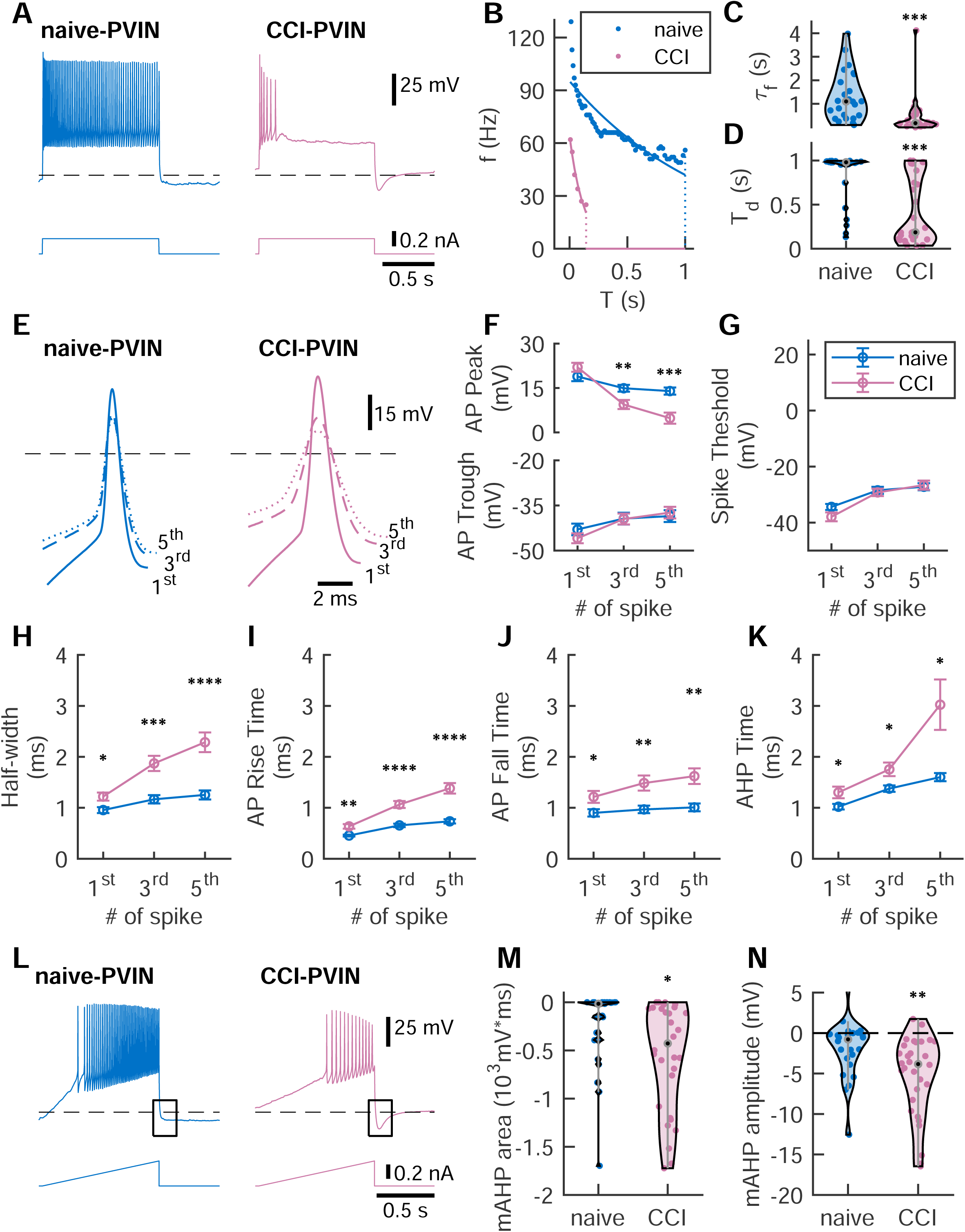
PVINs from chronic constriction injured mice show reduced excitability. **A**, Examples of electrophysiological responses to a 1-s long 200 pA step current of a naïve-PVIN (blue) and a CCI-PVIN (pink). The black dashed horizontal lines are at -70 mV. **B**, Instantaneous spike frequencies (dots) of the naïve-PVIN (blue) and CCI-PVIN (pink) in **A** measured as the inverse of inter-spike intervals and fitted to a single exponential decay function (lines), where the time constant *τ*_*f*_ refers to adaptation rate (**C**). For *τ*_*f*_: naïve, 1.31±0.21 s, n=24; CCI, 0.42±0.14 s, n=30; unpaired two-tailed t test, p=0.0007. Higher firing frequency adaptation rates of CCI-PVINs indicated their faster spike frequency adaptation. CCI-PVINs also had shorter discharge duration T_d_ (**D**). For T_d_: naïve, 0.82±0.06 s, n=24; CCI, 0.43±0.07 s, n=30; unpaired two-tailed t test, p=0.0001. **E**, Examples of the 1^st^ (solid line), 3^rd^ (dashed line) and 5^th^ (dotted line) APs from the spike trains triggered by a 1-s long 200 pA step current of a naïve-PVIN (blue) and a CCI-PVIN (pink). The black dashed horizontal lines are at 0 mV. Compared to the naïve condition, the peak amplitudes of 3^rd^ and 5^th^ APs were significantly decreased in CCI-PVINs (**F, top panel**) while the amplitudes of AP trough remained similar (**F, bottom panel**); the former was not due to differences in spike thresholds (**G**). PVINs generated wider APs as the spike number increased; this effect was stronger following CCI (**H**). In fact, the prolongation of the single spike duration was expressed in all phases of AP, including depolarizing (**I**), repolarizing (**J**) and after-hyperpolarization (AHP, **K**) phases. **F-K**, Error bars represent the SEM. Unpaired Student’s t test, *p<0.1, **p<0.01, ***p<0.001, ****p<0.0001. The values and statistical analysis are summarized in **Table 4**. **L**, Examples of electrophysiological responses to a 1-s long ramp current with a ramp rate of 0.4 pA/ms of a naïve-PVIN (blue) and a CCI-PVIN (pink). The black dashed horizontal lines are at - 70 mV. The mAHP was measured from the voltage trace 200 ms after the test stimulus (black boxes). The mAHP area (**M**) and mAHP amplitude (**N**) were significantly larger in CCI-PVINs. For mAHP area: naïve, -261.92±85.49 mV/ms, n=24; CCI, -558.11±102.59 mV/ms, n=30; unpaired two-tailed t test, p=0.036. For mAHP amplitude: naïve, -1.98±0.73 mV, n=24; CCI, - 5.25±0.91 mV, n=30; unpaired two-tailed t test, p=0.0092. (The voltages are not corrected for liquid junction potential in this figure and in Figs. 3-5)

The rapid spike frequency decrease of PVINs in nerve injured mice could result from the direct slowdown of the kinetics of individual action potentials (APs). During the 1-s step current stimulation, the shape of the action potential became smaller and wider as the spike number increased in both naïve and CCI conditions; however, this change was more drastic in CCI-PVINs (**Fig. 1E**). Indeed, the AP half-width increased significantly more in CCI-PVINs compared to naïve- PVINs (for the 1^st^, 3^rd^, and 5^th^ spikes, p=0.0144, 0.0004, and 3e-5, respectively, by unpaired Student’s t test; n=24 for naïve-PVINs and n=30 for CCI-PVINs; **Fig. 1H**). The prolonged spike durations of CCI-PVINs were further demonstrated by the increased duration of AP rise (upstroke), fall (downstroke) and after-hyperpolarization (for the 1^st^, 3^rd^, and 5^th^ spikes, AP rise: p=0.0015, 9e- 6, and 1e-6; AP fall: p=0.0377, 0.0058, and 0.0014; after-hyperpolarization: p= 0.0474, 0.0250, and 0.0132; by unpaired Student’s t test; n=24 for naïve-PVINs and n=30 for CCI-PVINs; **Fig. 1I-K**). The peak AP amplitude of CCI-PVINs was unaffected at the 1^st^ AP, but significantly decreased for the following APs (3^rd^ and 5^th^) (for the 1^st^, 3^rd^, and 5^th^ spikes, p=0.1702, 0.0097, and 0.0003, respectively, by unpaired Student’s t test; n=24 for naïve-PVINs and n=30 for CCI-PVINs; **Fig. 1F, top panel**). The minimal voltages at the after-hyperpolarization phase (i.e., AP trough), on the other hand, were not affected despite the longer duration of repolarization and AHP phases (for the 1^st^, 3^rd^, and 5^th^ spikes, p=0.2579, 0.9611, and 0.6840, respectively, by unpaired Student’s t test; n=24 for naïve-PVINs and n=30 for CCI-PVINs; **Fig. 1F, bottom panel**). These observations suggest that certain current(s) may be incrementally suppressing spike generation in PVINs, and their contribution is amplified after nerve injury. The spike thresholds were not different in the two conditions (for the 1^st^, 3^rd^, and 5^th^ spikes, p=0.0871, 0.7014, and 0.7903, respectively, by unpaired Student’s t test; n=24 for naïve-PVINs and n=30 for CCI-PVINs; **Fig. 1G**), indicating that major depolarizing currents, such as sodium currents, may not be affected.

After nerve injury, many PVINs showed a distinct medium after-hyperpolarization (mAHP) within 200 ms following a ramp current stimulus (mAHP amplitude: p=0.0092; mAHP area: p=0.0365; by unpaired Student’s t test; n=24 for naïve-PVINs and n=30 for CCI-PVINs; **Fig. 1L-N**), which is modulated by subthreshold conductances. These currents, such as calcium-activated potassium currents, are also implicated in spike frequency adaptation (Madison and Nicoll, 1984; Velumian and Carlen, 1999; Benda and Herz, 2003) and may be involved in regulating spike generation of PVINs.

It is important to note that we also observed a high prevalence of a voltage sag in recordings from both naïve- and CCI-PVINs. This sag is mediated by the activation of a hyperpolarization-activated cation current (*I*_ℎ_). However, no difference was observed between the two conditions (**Table 4**). While the *I*_ℎ_ has been previously implicated in spike frequency adaptation (Hewitt et al., 2021), it can also change the passive membrane properties of neurons including resting membrane potential and input resistance (Lupica et al., 2001). Evaluating both passive properties in naïve- and CCI- PVINs revealed that they are similar in two conditions (p>0.5 by unpaired Student’s t test; n=24 for naïve-PVINs and n=30 for CCI-PVINs; **Table 4**), suggesting that *I*_ℎ_ may not be the major current source that suppresses spike generation in PVINs after CCI.

### Computational modeling of PVIN firing activity predicts increased contribution of an SK-type conductance after nerve injury

To explore the ionic mechanisms controlling the electrical activity of PVINs following nerve injury, we employed a single-compartment HH type model of tonic-firing PVINs based on Bischop, et. al. 2012. The model was reparametrized to accurately capture the cycles of APs in the phase plane (V, dV/dt) generated by naïve-PVINs recordings (**Fig. 2A, Extended Fig. 2-1 & 2-2**). The model also reproduced the spiking behaviors of the neurons (referred to hereafter as naïve-PVIN model). Using this naïve-PVIN model, we sought to identify the minimal changes in ion channel expression required to produce the spike frequency adaptation observed in CCI-PVINs (**Fig. 2B-C**). We examined frequency adaptation properties as we varied the conductance density of each channel (**Fig. 2B-C**).

**Figure 2.**
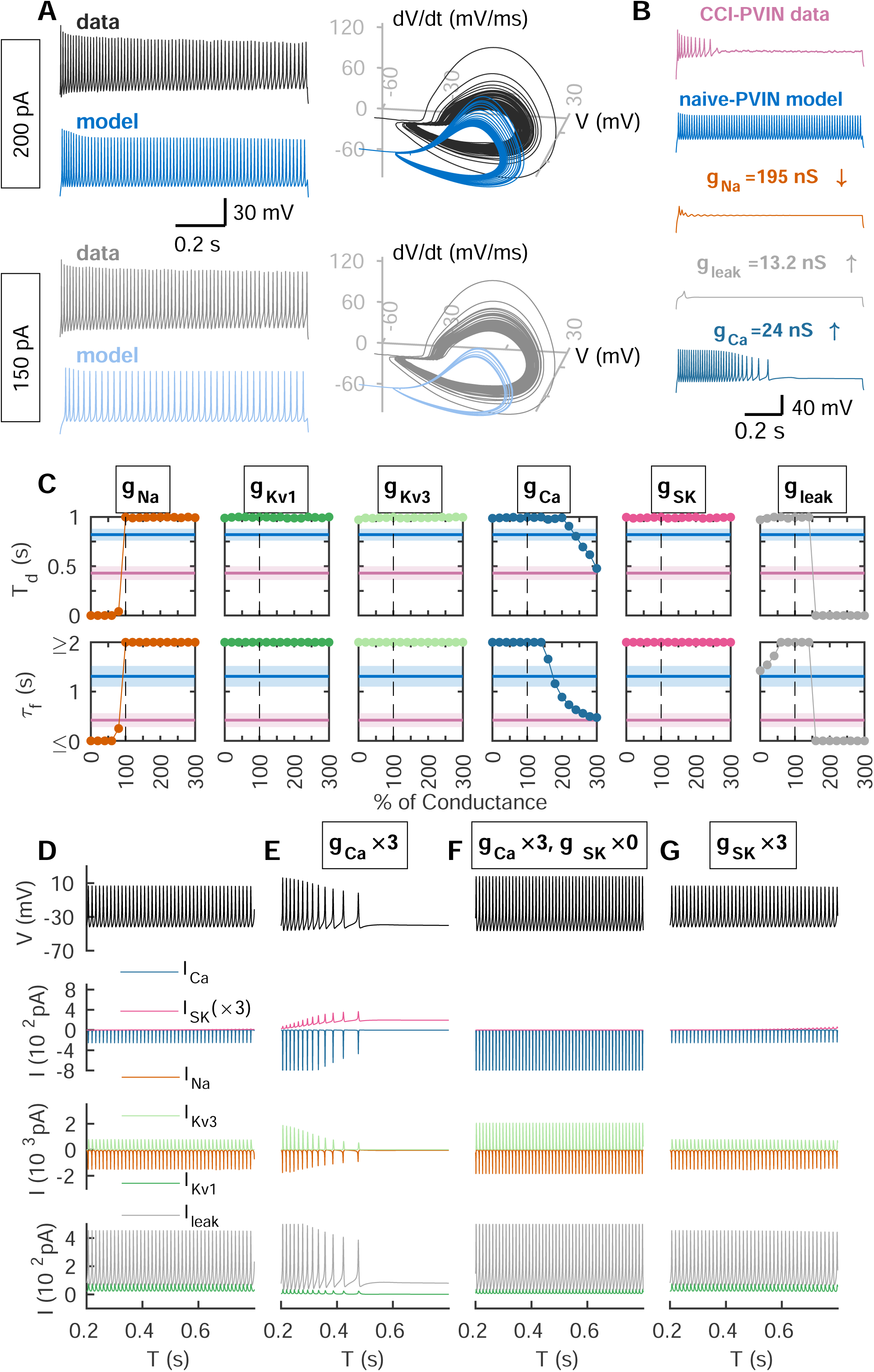
A Hodgkin-Huxley type model reproduces the excitability of naïve-PVINs. **A**, Simulated recordings from the model ([*B*_*tot*_]_*i*_ = 90 *μM*, dark and light blue) compared to naïve-PVIN recordings (black and grey) in response to multiple step currents (left panels). AP cycles of the same simulated and experimental recordings overlaid on top of each other in a 3D plot to facilitate comparison (right panels). **B**, Representative recording of CCI-PVIN (top, pink) along with simulated recordings of naïve-PVIN model with varied channel conductances as indicated in the legend (bottom recordings). **C**, Changes in spike frequency adaptation as the density of each ion channel conductance in naïve-PVIN model was varied. Line and colored regions in pink and blue are mean±SEM of discharge time T_d_ and frequency decay rate *τf* from naïve-PVINs (blue) and CCI-PVINs (pink). We compared how varying the maximal conductances of g_Na_ (orange), g_Kv1_ (dark green), g_Kv3_ (light green), g_Ca_ (blue-green), g_SK_ (magenta), or g_leak_ (grey) alter the activity of naïve-PVIN model to that comparable to CCI-PVIN data (pink regions). **D-G**, Simulated ionic currents of naïve-PVIN model (**D**), the naïve model with 3-fold increase in g_Ca_ (**E**), the naïve model with 3-fold increase in g_Ca_ and a decrease to zero in g_SK_ (**F**), and the naïve model with 3-fold increase in g_SK_ (**G**) in response to a 0.8-s long 100 pA current. Three-fold increase in g_Ca_ causes a large increase in I_Ca_ (blue-green) and I_SK_ (magenta), while the other ionic currents in the model, I_Na_ (orange), I_Kv1_ (dark green), I_Kv3_ (light green), and I_leak_ (grey) are negligibly affected (**D**).

A decrease of more than 20% in the sodium channel conductance density was sufficient to silence the neuron (**Fig. 2B, orange trace**). As for leak conductance density, increasing it moderately (by 1.6-fold) did not affect firing activity, but a further increase (>1.8-fold) significantly diminished the firing activity (**Fig. 2B, grey trace**). However, the loss in the firing activity of PVINs under these two conditions led to a substantial reduction in the 1st-spike peak amplitude, which is not observed experimentally in CCI-PVINs recordings. This suggests that these conductances are not responsible for the change in firing activity of PVINs after nerve injury.

On the other hand, increasing the calcium conductance density by 3-fold led to a frequency decay comparable to the data of CCI-PVINs (**Fig. 2B, blue-green trace; Fig. 2C, blue-green trace**). Calcium influx through high voltage-gated calcium channels favor spike frequency adaptation via the coupled activation of small-conductance calcium-activated potassium (SK) channels (Hallworth et al., 2003). Indeed, compared to the control simulation (**Fig. 2D**), increasing the calcium conductance density 3-fold produced larger I_SK_ (**Fig. 2D-E, pink traces**) as I_Ca_ increases (**Fig. 2D- E, blue-green traces**). This model exhibited spike frequency adaptation that can be reverted to tonic firing by blocking SK conductance (**Fig. 2F**). Given the evidence that SK channels also mediate the mAHP (Sah and Faber, 2002), we hypothesized that increasing SK conductance density would oppose the tonic firing activity of PVINs, similar to what is seen experimentally after nerve injury. However, a 3-fold increase in the maximal ionic conductance of SK channels (**Fig. 2G**) did not reproduce the spike frequency adaptation seen in PVINs after nerve injury. This suggests that although SK channels remain a good candidate to explain frequency adaptation in CCI-PVINs, additional factors must be involved. To determine whether a greater increase in SK channel conductance density was necessary to reproduce the CCI-PVIN observation, we increased g_sk_ more than 10-fold (**Fig. 3A-B**). Interestingly, even with such unphysiological levels, the firing activity of PVINs only showed modest adaptation while the discharge duration did not change (**Fig. 3A-B**).

**Figure 3.**
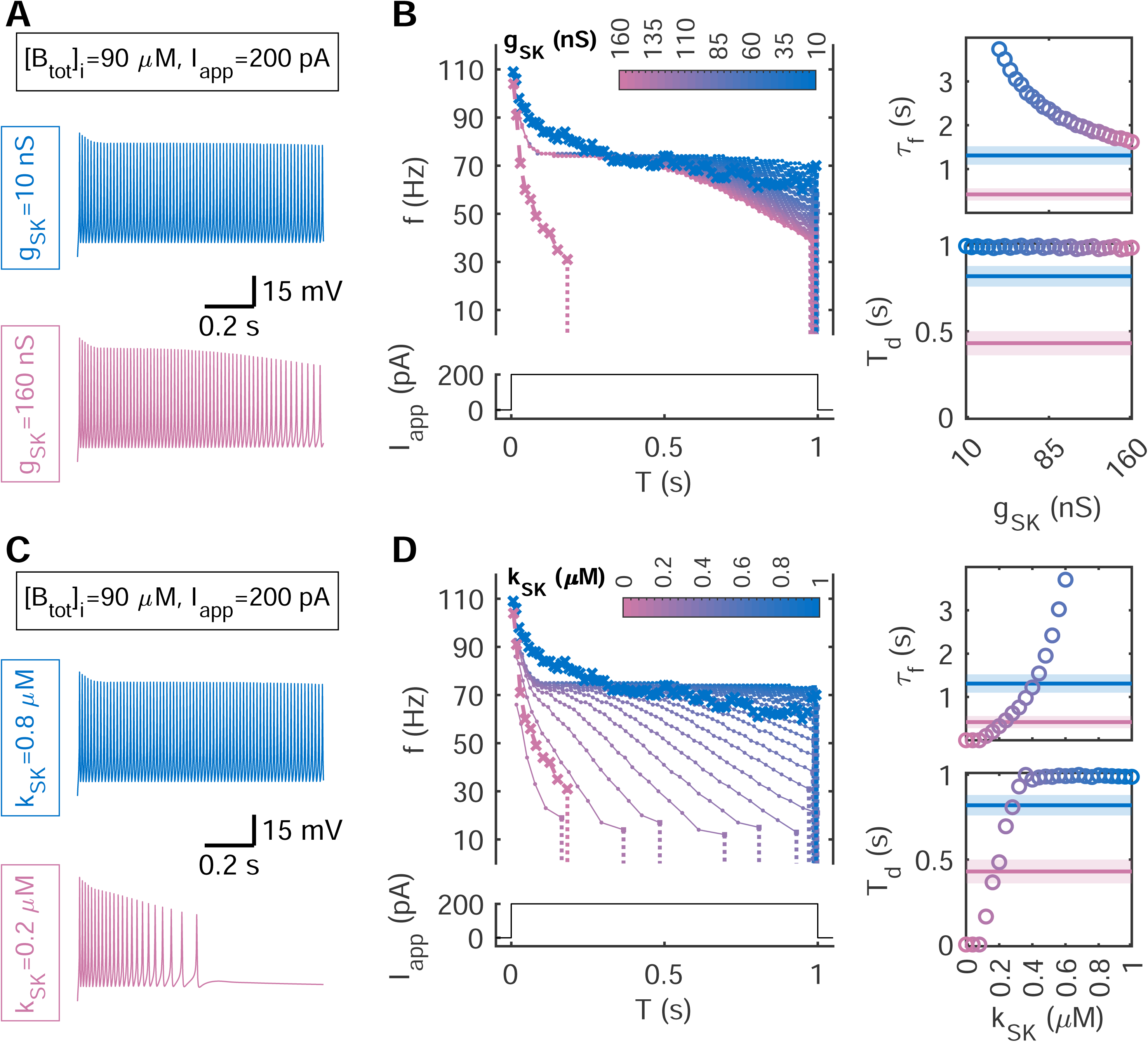
Computational and pharmacological approaches implicate SK channels in inducing spike frequency adaptation in CCI-PVINs. **A**, Simulated recordings from the naïve-PVIN model ([*Btot*]*i* = 90 *μM*) with g_SK_=10 nS (blue) and 160 nS (pink) in response to a 1-s long 200 pA step current. **B**, Comparison of instantaneous firing frequencies of simulated recordings (dotted lines) from the naïve-PVIN model ([*Btot*]*i* = 90 *μM*) with varied values of g_SK_ and of electrophysiological recordings from a naïve-PVIN (blue crossed lines) and a CCI-PVIN (pink crossed lines). For the simulated model (dotted lines), the value of g_SK_ is in the range between 10 nS (blue) and 160 nS (pink). The right panels are the quantifications of frequency decay rate *τ*_*f*_ (top-right panel) and discharge time T_d_ (bottom-right panel). Lines and colored regions in pink and blue are mean±SEM from naïve-PVINs (blue) and CCI-PVINs (pink). **C**, Simulated recordings from the naïve-PVIN model ([*Btot*]*i* = 90 *μM*) with k_SK_=0.8 *μ*M (blue) and 0.2 *μ*M (pink) in response to a 1-s long 200 pA step current. **D**, Same as B, except that we varied the parameter k_SK_, representing the calcium-concentration-level for half-maximal activation of SK channels, between 0 *μ*M (pink) and 1 *μ*M (blue).

Another parameter determining the gating of SK channels is the calcium sensitivity. The calcium sensitivity of the SK channels has been reported to be around 0.7 μM under control conditions (Hirschberg et al., 1998) and it is under physiological regulation such as the intracellular calcium dynamics (Hallworth et al., 2003; Fakler and Adelman, 2008) and phosphorylation (Bildl et al., 2004; Allen et al., 2007; Zhang et al., 2014; Mizukami et al., 2015). We therefore examined whether a change in calcium sensitivity could explain the increased contribution of SK channels in CCI-PVINs. Our results indicate that an increase in calcium sensitivity of SK channels *k*_*SK*_ to ≤0.2 μM can effectively alter the discharge time and frequency decay rate in a manner similar to CCI-PVINs (**Fig. 3C-D**). This was confirmed experimentally by pharmacological manipulation of SK channels (**Fig. 4**). We hypothesized that activating SK channels in a naïve-PVIN would cause a transition in the firing pattern from tonic to adaptive. The SK channel agonist 1-EBIO can cause a leftward-shift in the half-maximal activation curve of SK channels to *k*_*SK*_ =0.1 *μM* (Pedarzani, et al., 2001). When 1-EBIO (300 μM) was tested against tonic firing naïve-PVINs, we observed the development of a clear spike frequency adaptation in naïve-PVINs (n=5; p<0.5 by paired t test; **Fig. 4** and **Table 5**), which is consistent with the simulations. Therefore, both computational and experimental evidence demonstrated the role of SK channels in modulating the excitability of PVINs following nerve injury.

**Figure 4.**
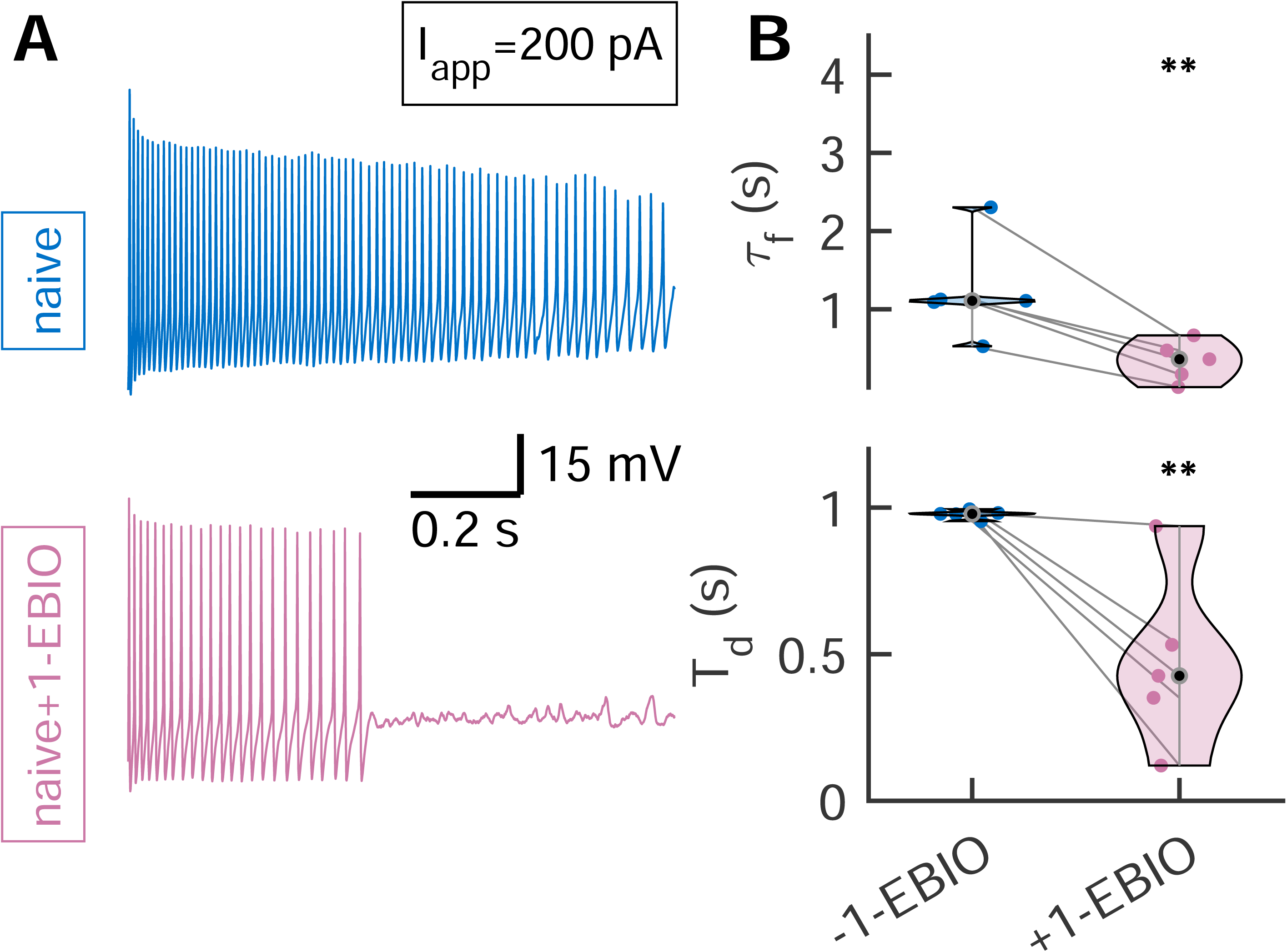
**A**, Examples of electrophysiological responses to a 1-s long 200 pA step current of a naïve-PVIN before (blue) and after the application of 1-EBIO (pink). **B-C**, Quantification of 1- EBIO’s impact on the electrical responses of persistent firing naïve-PVINs including frequency decay rate *τf* (**B, top panel**) and discharge time T_d_ (**B, bottom panel**). For *τf*: naïve, 0.95±0.27 s; naïve+1-EBIO, 0.25±0.10 mV/ms; p=0.0050; for T_d_: naïve, 0.76±0.14 s; naïve+1-EBIO, 0.35±0.12 s; p=0.0046; paired Student’s t test; n=5; **p<0.01.

**Table 5.** Electrophysiological features of tonic firing naïve-PVINs (n=5) before and after the application of 1-EBIO. Data are mean±SEM.

| Properties | - 1-EBIO | +1-EBIO | p value |
| --- | --- | --- | --- |
| Resting membrane potential (mV) | -49.96±0.94 | -52.44±3.45 | 0.4407 |
| Input resistance (MΩ) | 243.66±23.22 | 217.41±17.05 | 0.1336 |
| Rheobase (pA) | 18.43±1.87 | 29.67±9.28 | 0.3246 |

### A decrease in cytosolic calcium buffer is sufficient to reproduce spike frequency adaptation observed in CCI-PVINs

We next investigated the potential mechanisms that could lead to the potentiation of SK channels in CCI-PVINs. We hypothesized that the hyperactivation of SK channels is induced by the dysregulation of intracellular calcium dynamics, to be specific, the depletion of the cytosolic calcium buffer. According to this framework, the large calcium influx elicited by neuronal spiking escapes chelation by calcium buffer and triggers the opening of SK channels, thus leading to spike frequency adaptation. By simulating the naïve-PVIN model after decreasing its intracellular concentration of calcium buffer ([B_tot_]_i_) by 80 μM, we obtained voltage responses and spike shapes comparable to the CCI-PVIN recordings (**Fig. 5A**). Indeed, the model showed that in response to a depolarizing current, the initial fast spiking response drives free calcium into the cytoplasm, a great portion of which fails to be chelated due to the limited amount of calcium buffer. As a consequence, calcium triggers a large potassium efflux through SK channels until the increasing hyperpolarizing power silences the cell (**Fig. 5B**).

**Figure 5.**
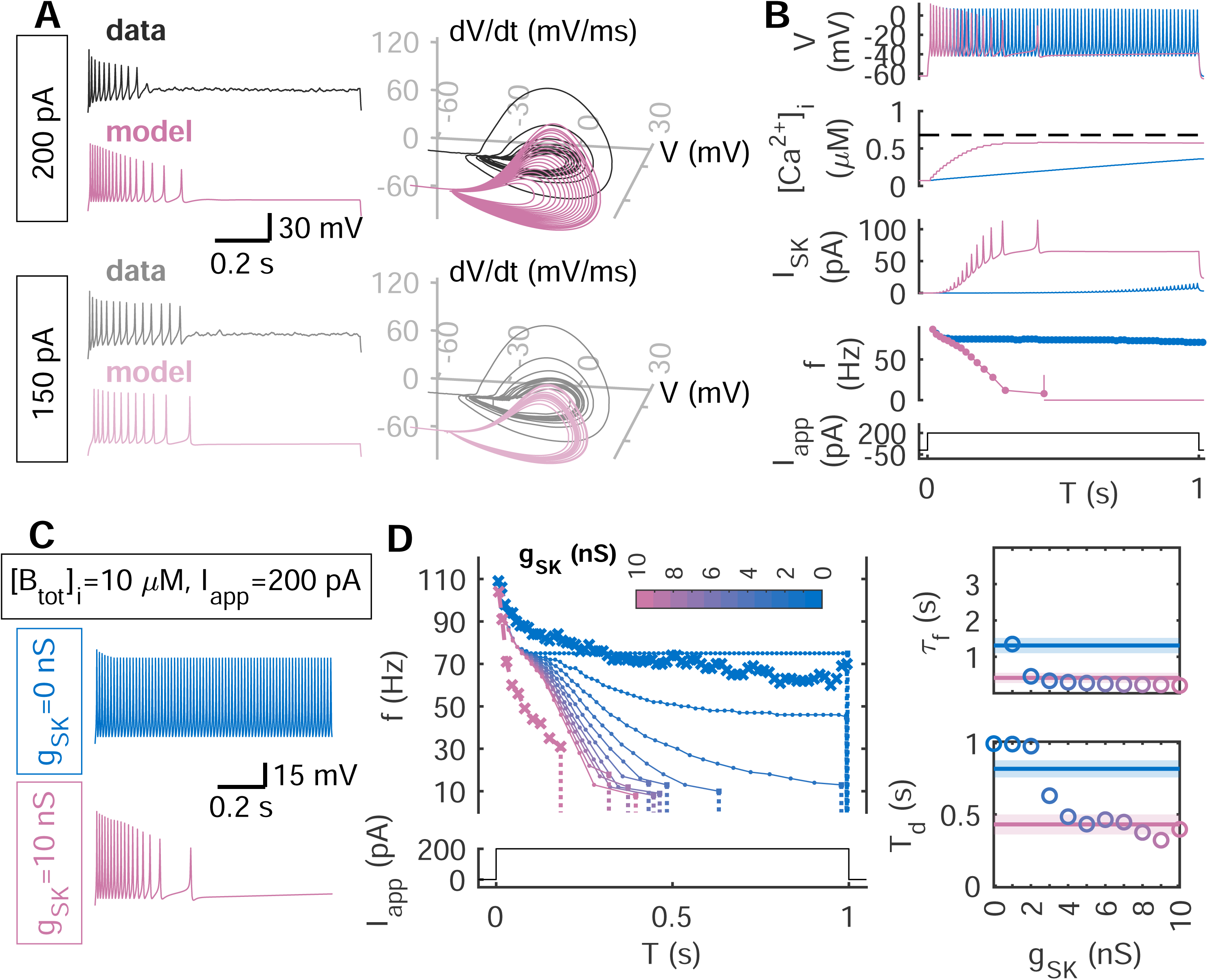
Calcium buffer depletion in naïve-PVIN model reproduces the firing behavior of CCI- PVINs. **A**, Simulated recordings from the PVIN model with reduced buffer concentration ([*B*_*tot*_]_*i*_ = 10 *μM*, dark and light pink) compared to CCI-PVIN recordings (black and grey) in response to multiple step currents (left panels). AP cycles of the same simulated and experimental recordings overlaid on top of each other to facilitate comparison (right panels). **B**, The model showed that, compared to naïve-PVIN model (blue), less calcium buffer ([*B*_*tot*_]_*i*_ = 10 *μM*, pink) allows fast and large accumulation of [*Ca*^2+^]_*i*_ which in turn hyper-activates SK channels and ceases firing. The black dashed line in the second panel from the top is at 0.8 *μM*, which refers to the calcium-concentration-level for half-maximal activation of SK channels. **C**, Simulated recordings from the CCI-PVIN model ([*Btot*]*i* = 10 *μM*) with g_SK_=0 nS (blue) and 10 nS (pink) in response to a 1-s long 200 pA step current. **D**, Comparison of instantaneous firing frequencies of simulated recordings (dotted lines) from the CCI-PVIN model ([*B*_*tot*_]_*i*_ = 10 *μM*) with varied values of g_SK_ and of electrophysiological recordings from a naïve-PVIN (blue crossed lines) and a CCI-PVINs (pink crossed lines). For the simulated model (dotted lines), the value of g_SK_ is in the range between 0 nS (blue) and 10 nS (pink). The right panels are the quantifications of frequency decay rate *τf* (top-right panel) and discharge time T_d_ (bottom-right panel). Lines and colored regions in pink and blue are mean±SEM from naïve-PVINs (blue) and CCI-PVINs (pink).

The model further predicted that in conditions where the calcium buffer levels are low, blocking SK channels can rescue tonic firing in CCI-PVINs (**Fig. 5C-D**). This was done by modifying the maximal channel conductance *g*_*SK*_ to 0 nS. Consistent to the simulation, the SK channels blocker apamin (200 nM) applied to CCI-PVINs significantly rescued the spike frequency adaptation (n=9; p<0.5 by paired t test; **Fig. 6** and **Table 6**) to that of naïve-PVINs level.

**Figure 6.**
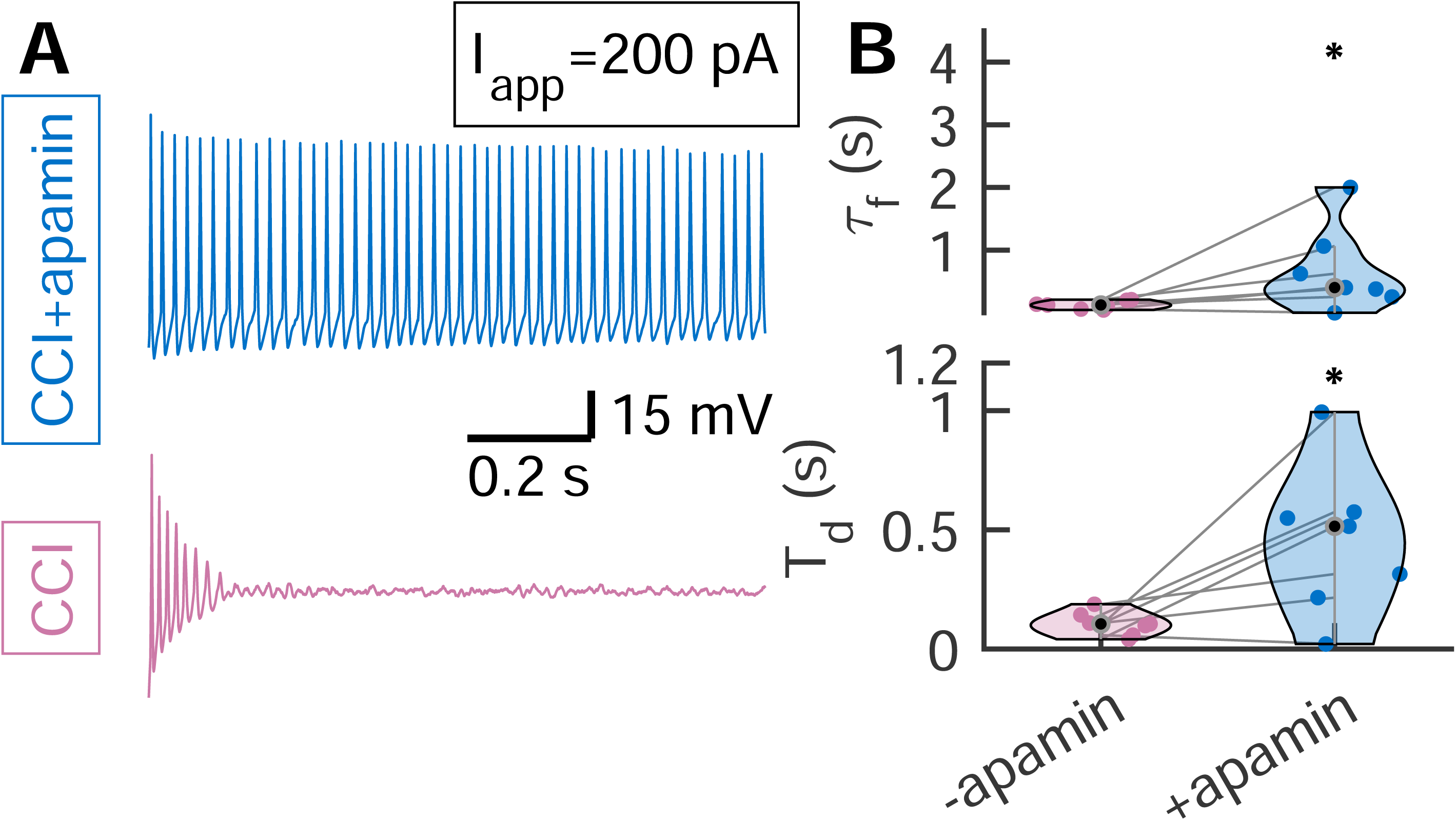
**A**, Examples of electrophysiological responses to a 1-s long 200 pA step current of a CCI-PVIN before (pink) and after the application of apamin (blue). **B-C**, Quantification of apamin’s impact on the electrical responses of transient firing CCI-PVINs including frequency decay rate *τf* (**B, top panel**) and discharge time T_d_ (**B, bottom panel**). For *τf*: naïve, 0.13±0.02 s; naïve+1-EBIO, 0.67±0.25 mV/ms; p=0.0314; for T_d_: naïve, 0.11±0.02 s; naïve+1-EBIO, 0.45±0.12 s; p=0.0128; paired Student’s t test; n=7; *p<0.05.

**Table 6.** Electrophysiological features of transient firing CCI-PVINs (n=7) before and after the application of apamin. Data are mean±SEM.

| Properties | -apamin | +apamin | p value |
| --- | --- | --- | --- |
| Resting membrane potential (mV) | -52.86±1.58 | -56.93±3.59 | 0.2324 |
| Input resistance (MΩ) | 210.55±28.94 | 195.40±23.83 | 0.4672 |
| Rheobase (pA) | 32.43±5.78 | 35.61±6.13 | 0.7550 |

### Slow-fast analysis of the HH model revealed calcium-dependent dynamics underlying CCI- PVIN state transitions from spiking to quiescence

Under stable current stimulation, CCI-PVINs predominantly exhibit transient firing: upon a depolarizing current injection, the neuron initiates spiking and transits to a quiescent state. Upon examining spike-amplitude adaptation rate during this transient firing, we identified 2 distinct patterns of state transitions in CCI-PVINs (n=30): *on-off switching* and *damped oscillation* (**Fig. 7A**). The on-off switching refers to the class of CCI-PVINs displaying repetitive suprathreshold spiking that eventually switches abruptly to quiescent state (**Fig. 7A, top panel**). This preferentially occurred following modest current injections (*I*_*app*_ = 150 *pA*, **Fig. 7B, top panel**). In contrast, the damped oscillation refers to the class of CCI-PVINs that display APs whose amplitudes gradually decrease to subthreshold level before reaching the quiescent state (**Fig. 7A, bottom panel**). This latter case was more likely to be observed for higher current injections (*I*_*app*_ = 200 *pA*, **Fig. 7B, bottom panel**).

**Figure 7.**
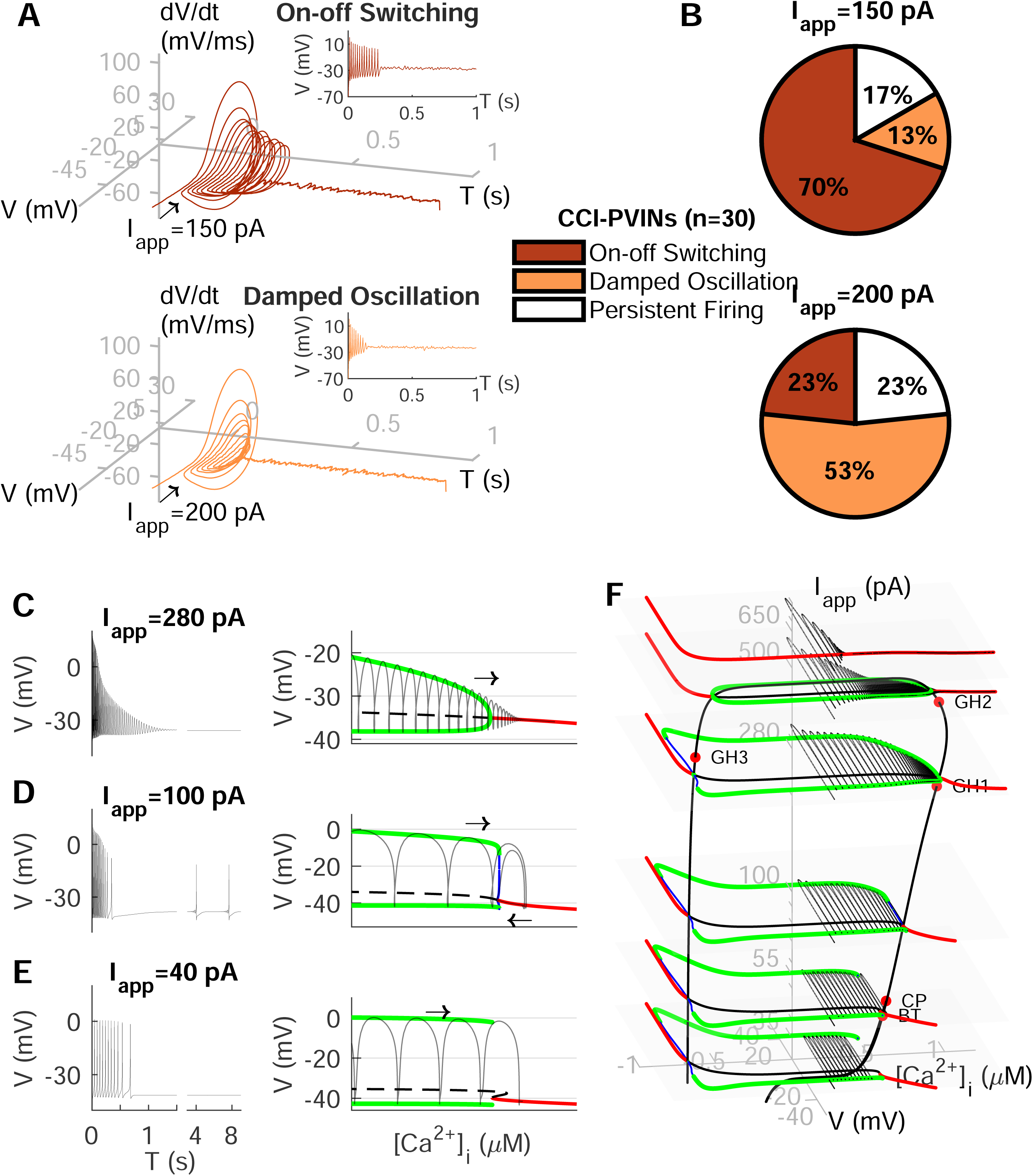
Bifurcation analysis reveals that intracellular calcium controls the dynamics underlying state transitions of CCI-PVINs. **A**, Examples of temporal evolution of AP cycles and the voltage trace (**insets**) produced by a CCI-PVIN that exhibit the state transition type “on-off switching” (dark red, **top panel**) at I_app_=150 pA and “damped oscillation” (orange, **bottom panel**) at I_app_=150 pA. **B**, Firing pattern distribution of CCI-PVINs (n=30) triggered by a 1-s long 150 pA (**top panel**) or a 200 pA (**bottom panel**) step current. The on-off switching pattern prevalently occurred at *I*_*app*_=150 pA, while the damped oscillation pattern were more likely to occur at higher injection currents. **C-E**, Examples of CCI-PVIN model simulations (**left panels**) with their corresponding one-parameter bifurcation diagrams with respect to [*Ca*^2+^]_*i*_ (**right panels**) in response to these injection currents: 280 pA (**C**), 100 pA (**D**) and 40 pA (**E**). In the bifurcation diagrams, black dashed lines represent branches of unstable steady states, red solid lines represent branches of stable steady states, green (blue) lines represent envelopes of stable (unstable) periodic orbits, black solid lines represent the voltage simulations superimposed on the one-parameter bifurcations, and the rightward arrows refer to the direction of the trajectories. The periodic envelopes in these diagrams emanate from either supercritical Hopf (top) subcritical Hopf (middle) or saddle-node on an invariant circle SNIC (SNIC, bottom) bifurcations. Notice that in **D** and **E**, additional delayed spikes were generated through an elliptic and parabolic type firing mechanism, respectively. **F**, The evolution of one-parameter bifurcation diagrams with respect to [*Ca*^2+^]_*i*_ as the injection current increases (from bottom to top, *I*_*app*_=35, 55, 100, 280, 500, 650 *pA*); these bifurcation diagrams were stacked on top of each other to demonstrate how a two-parameter bifurcation diagram can be formed.

Such state transitions correspond to “bifurcation” from a dynamical system’s point of view. To examine the dynamical mechanisms underlying these state transitions, we performed slow-fast decomposition analysis. In this analysis, the variable that changed most slowly, compared to other variables, was treated as a “parameter”. In our model, cytosolic calcium concentration [*Ca*^2+^]_*i*_ was found to be the slowest variable and as such was used as a bifurcation parameter to determine how it dictates the switching between the two states of the model: spiking and quiescence. To do so, we performed a one-parameter bifurcation analysis, which plots the steady state values of the membrane voltage *V* as a function of the new parameter [*Ca*^2+^]_*i*_ at various values of the injected current *I*_*app*_ (**Fig. 7C-F**).

According to the model, when *I*_*app*_ is lower than ∼30 pA, the voltage is quiescent, i.e., non-spiking. At *I*_*app*_ = 40 pA, the model exhibits periodic spiking that switches to quiescence via a saddle-node on invariant circle (SNIC) bifurcation (**Fig. 7E, right panel**). Superimposing the voltage trace on this one-parameter bifurcation diagram (**Fig. 7E, right panel**) showed that this curve initially oscillates between the two envelopes of stable limit cycles until it crosses the SNIC bifurcation and settles at the stable branch of equilibria. This type of behavior is consistent with “subHopf/circle” bursting (Izhikevich, 2000). Because calcium recovery rate (denoted by *γ* in the PVIN model, see Materials and methods) is slow, the next spike following the quiescent state occurs at a time that is rather late (∼5 s after initial spiking) and is likely absent in the 1-s long recordings. Interestingly, increasing *I*_*app*_ to 100 pA causes the limit points (LPs) to disappear. Superimposing the voltage trace on this one-parameter bifurcation diagram (**Fig. 7E**) revealed that the transition to resting state occurs via a fold limit cycle, while the transition to spiking is via a subcritical Hopf bifurcation. This produced the activity that is consistent with an “elliptic” type (also called “subHopf/fold cycle”) bursting (Izhikevich, 2000). In these scenarios (“subHopf/circle” bursting and “subHopf/fold cycle”), the model exhibits state transitions corresponding to the “on-off switching” transition observed in the data. For higher injected current (*I*_*app*_ = 280 *pA*, **Fig. 7C**), the oscillations in the superimposed voltage trace disappeared via supercritical Hopf bifurcation, producing outcomes consistent with the “damped oscillation” pattern. It is important to note here that further increasing *I*_*app*_ would make the model eventually non-responsive due to a depolarization block.

### The two-parameter bifurcation analysis revealed that a generalized Hopf bifurcation underlies the transition between the two patterns of CCI-PVINs as the injected current increases

The relationship between the two patterns of CCI-PVINs and the *I*_*app*_ can be elucidated using a two-parameter bifurcation analysis in the ([*Ca*^2+^]_*i*_, *I*_*app*_)-plane (**Fig. 7F and Fig. 8**). This parameter space can be divided by the one-parameter bifurcation curves of the Hopf bifurcation point (HB) and the LP into 3 regions: the region of stable equilibrium (yellow), the region of unstable equilibrium (white) and the region of stable periodic oscillations bounded to the right by the Hopf bifurcation (green). The two-parameter bifurcation points, obtained from the one-parameter bifurcation curves, further partitioned the ([*Ca*^2+^]_*i*_, *I*_*app*_)-plane into regions of transition patterns corresponding to those observed in CCI-PVIN recordings (bidirectional arrows in **Fig. 8**). Importantly, the two-parameter bifurcation analysis revealed that, as *I*_*app*_ increases, a generalized Hopf bifurcation point (GH) represents the condition when the one-parameter HB changes from subcritical to supercritical, or electrophysiologically, when the firing pattern switches from on-off switching to damped oscillation. Taken together, these results indicate that the slow dynamics of free cytosolic calcium plays a critical role in modulating the transient firing behavior in CCI-PVINs; it also highlights the ability of the model to reproduce and explain the transition in pattern from on-off switching to damped oscillation as *I*_*app*_ increases, in a manner similar to that seen in CCI-PVIN recordings.

**Figure 8.**
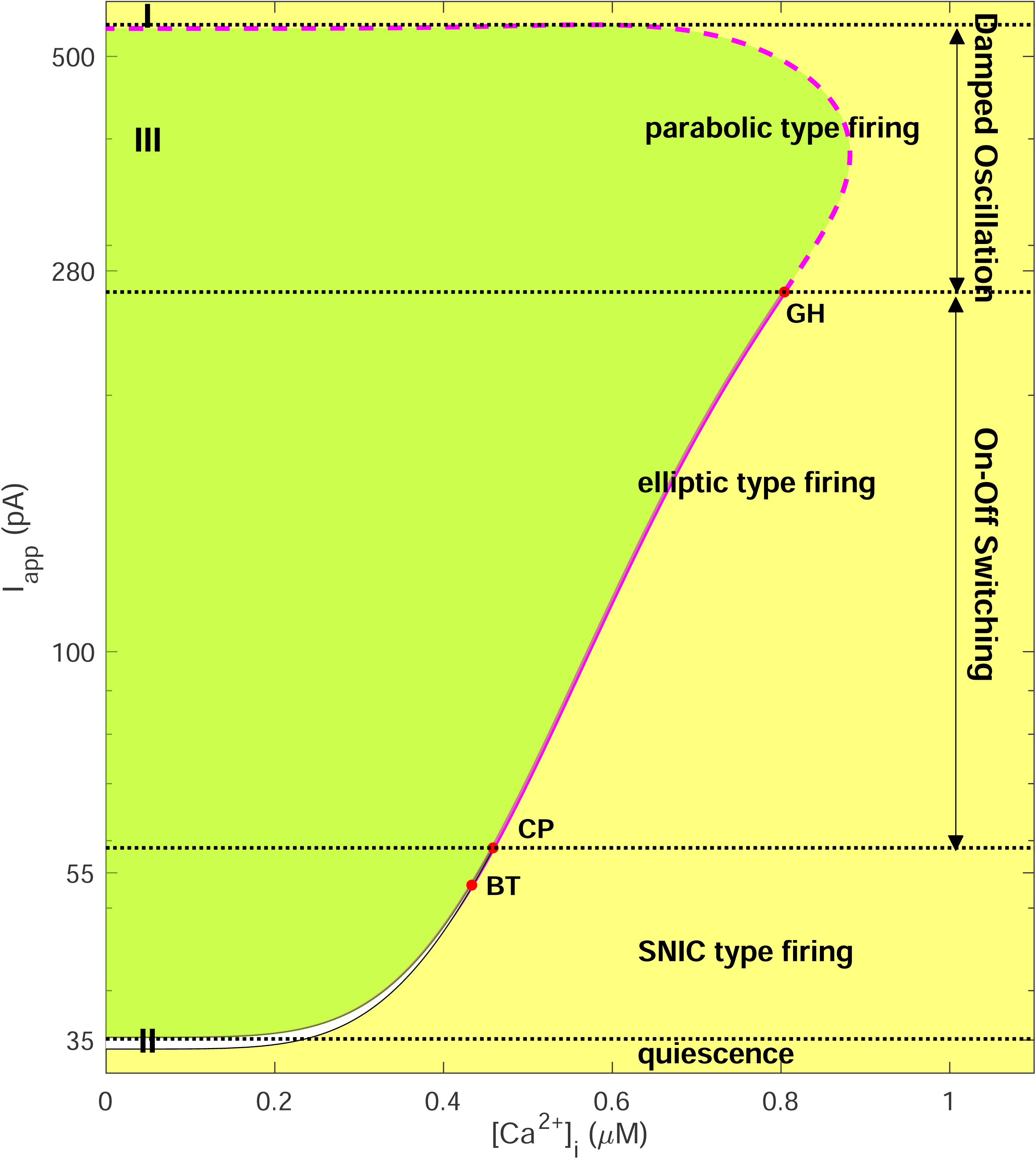
Color-map of the various regimes of behavior defined by the two-parameter bifurcation of the CCI-PVIN model in the ([*Ca*^2+^]_*i*_, *I*_*app*_) plane. Two-parameter continuation of the bifurcation points in Fig. 7F, obtained by varying the two parameters [*Ca*^2+^]_*i*_ and *I*_*app*_ ; this generated three distinct regimes of behavior demarcated by lines representing the subcritical Hopf bifurcation (solid magenta line), supercritical Hopf bifurcation (dashed magenta line) and SNIC bifurcation (black dash-dotted line). These regimes correspond to Region I of stable equilibrium points (yellow), Region II of unstable equilibrium points (white) and Region III of stable periodic orbits responsible for repetitive firing (green). Here, we only discuss the dynamics occurring at [*Ca*^2+^]_*i*_ ≥ 0 *μM* (the unphysiological relevant region [*Ca*^2+^]_*i*_ < 0 *μM* not shown). The two- parameter bifurcation points (red dots) are: GH--generalized Hopf bifurcation, CP--cusp bifurcation, BT--Bogdanov-Takens bifurcation.

### Calcium buffer-depleted PVIN model shows an impaired inhibitory control over PKCγ- expressing interneurons in a simulated Aβ fiber-driven neural circuit

The commonly used current-clamp protocols consist of injecting simple and smooth step or ramp waveforms of currents. These protocols have been used to standardize the comparison of firing behaviors among different types of neurons for many years. However, their stimulations trigger neural responses that may not well-represent how a neuron reacts and functions in the context of neural circuit, given that the realistic endogenous inputs are non-linear and correlated (Silver, 2010; Druckmann et al., 2011; Szabó et al., 2021). To examine the effects of a decrease in calcium buffer concentration on the firing activity of PVINs elicited by presynaptic input from the low-threshold A*β* fibers, we used a more realistic input current: a 3-sec Poisson-distributed presynaptic current (**Fig. 9A-C**). The frequency of the Aβ stimulus (f_Aβ_) was chosen to correlate with the mechanical force. When a light (*f*_*Aβ*_ = 5 *Hz*) or an intense (*f*_*Aβ*_ = 15 *Hz*) mechanical force (**Fig. 9A-C**) was applied, the PVIN model with reduced calcium buffer failed to actively respond in both scenarios compared to naïve-PVIN model; indeed, the input-output frequency ratio (PV/Aβ) decreased from 8.249 to 0.373 between naïve- and CCI-PVIN models, respectively. This result indicates that calcium buffer-depleted PVINs have a diminished ability for frequency encoding.

**Figure 9.**
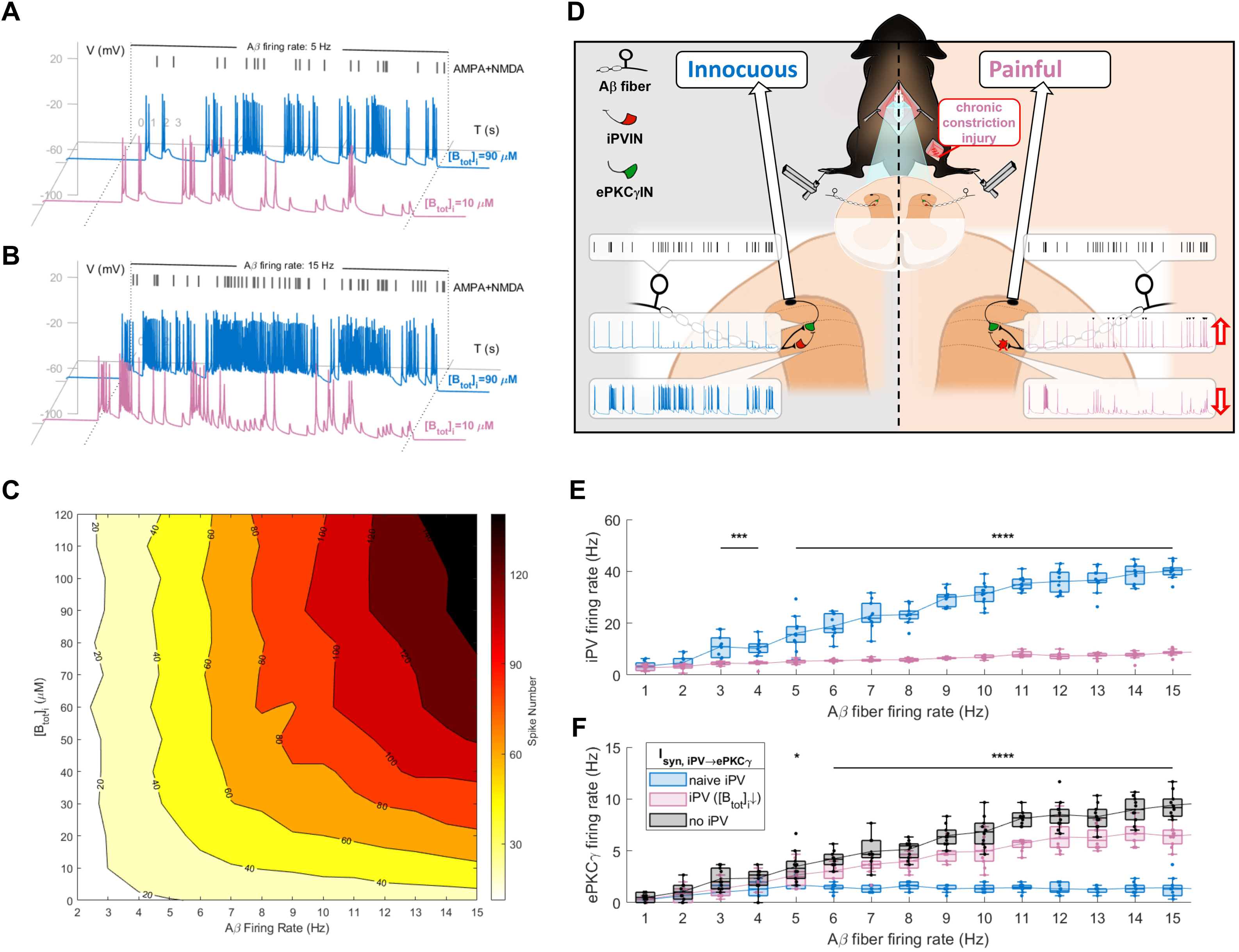
The impaired excitability of PVINs with reduced calcium buffer failed to cause sufficient inhibitory-control of *Aβ* fiber-mediated pain sensation pathway. **A**, Simulated time courses of the PVIN model with high ([*B*_*tot*_]_*i*_ = 90 *μM*, blue) and low ([*B*_*tot*_]_*i*_ = 10 *μM*, pink) calcium buffer concentration in response to 5 Hz synaptic inputs representing light mechanical stimulus. **B**, Same as in **A**, but using 15 Hz synaptic inputs representing intense mechanical stimulus that maximizes the firing rate of *Aβ* fibers. **C**, Heatmap illustrating the number of spikes generated by the PVIN model with varied [*B*_*tot*_]_*i*_ in response to different presynaptic firing rates. PVIN model of reduced [*B*_*tot*_]_*i*_ generated less spikes regardless of the presynaptic firing rate. **D**, Schematic of a simplified circuit representing the *Aβ* fiber-mediated pain sensation pathway in spinal dorsal horn of the mice in naïve (left panel) and CCI condition (right panel). Both inhibitory PVIN (green) and excitatory PKC*γ*IN (red) receive presynaptic inputs from the same *Aβ* fiber; the excitatory PKC*γ*IN also receive inhibitory presynaptic inputs from the PVIN. The calcium buffer depleted PVIN (right panel) failed to inhibit the *Aβ* fiber-triggered activity of PKC*γ* IN. The spikes generated by PKC*γ*IN that escaped PVIN’s inhibition was marked by black inverted triangles. **E-F**, The firing frequency of *PKCγ* IN and PVIN in response to varied frequencies of A *β* fiber stimulation. Reducing [*B*_*tot*_]_*i*_ from 90 *μM* (blue) to 10 *μM* (pink) in PVIN led to significant decrease in PVIN activities (**E**) and increase in PKC*γ*IN activities when the firing frequency of *Aβ* fiber is higher than 5 Hz (**F**). Black curve in **F** represents the scenario when the excitatory PKC*γ*IN receives only the excitatory presynaptic input from the *Aβ* fiber. Paired Student’s t test, *p<0.1, ***p<0.001, ****p<0.0001.

To further assess the role of PVINs on controlling electrical signaling in the spinal dorsal horn, we embedded the PVIN model in an “*in vivo*-like” neural circuit (**Fig. 9D**) that comprises both their pre- and post-synaptic components. Specifically, this circuit model included the Poisson-distributed excitatory presynaptic currents to represent the inputs from the A*β* fibers, as well as another HH type model describing the excitability of a PVIN post-synaptic target: the excitatory interneuron expressing protein kinase C gamma (PKC*γ*IN). The A*β* fiber-like presynaptic current was applied on both the inhibitory PVIN model and the excitatory PKC*γ*IN model, the latter of which also received inhibitory synaptic input from the PVIN model. The efficiency of signal transformation of this simplified neural circuit was expressed by the frequency input-output relationship (**Fig. 9E-F, Table 7**). Under normal condition, the firing frequency of PVIN model increased with that of the Aβ fiber increase (**Fig. 9E, blue trace**). This allowed the fast-spiking PVIN model to efficiently reduce Aβ-driven activation of PKC*γ*IN model to less than ∼5 Hz, irrespective of the firing rate of A*β* fiber input (*R*^2^=0.183) (**Fig. 9F, blue trace**). When the firing frequency of the PVIN model was significantly decreased due to a reduction in calcium buffer (**Fig. 9E, pink trace**), the PKC*γ*IN model was able to linearly mirror the frequency-coding from A*β* fibers at a rate of 2.26 (firing ratio of PKCγIN/Aβ) (*R*^2^ = 0.804) (**Fig. 9F, pink trace**). The importance of calcium buffer is further supported by the observation that removing PVINs altogether produced the same level of disinhibition (**Fig. 9F, black trace**). This disinhibition appeared at *f*_*Aβ*_ = 5 *Hz* and was more pronounced as *f*_*Aβ*_ increased. Taken together, these results further demonstrate the importance of calcium buffer in modulating the ability of PVINs to prevent the information output of A*β* fibers from activating dorsal horn pain pathways.

**Table 7.** The synaptic efficiencies on information transduction of PVIN model and PKCγ model.

| Properties | iPV |  | ePKCγ |  |  |
| --- | --- | --- | --- | --- | --- |
|  | naive | CCI | naïve iPV<br>([Ca<sup>2+</sup>]<sub>i</sub> =<br>90 μM) | CCI iPV<br>([Ca<sup>2+</sup>]<sub>i</sub> =<br>10 μM) | no iPV |
| Correlation Coefficient<br>(r) | 0.932 | 0.888 | 0.427 | 0.897 | 0.876 |
| Linear Regression:<br>Slope | 8.249 | 0.373 | 0.185 | 2.26 | 1.606 |
| Linear Regression:<br>R-square (R<sup>2</sup>) | 0.869 | 0.789 | 0.183 | 0.804 | 0.767 |

## Discussion

In the dorsal horn of the spinal cord, PVINs represent a critical source of inhibition that prevents touch inputs from activating pain pathways (Petitjean et al., 2015; Cao et al., 2022; Medlock et al., 2022). However, after nerve injury, it is believed that the output of these neurons is significantly decreased, leading to a mechanical allodynia, whereby gentle touch stimuli produce pain responses. This decreased output can be the result of at least three non-exclusive mechanisms: (1) a decrease in the excitatory drive that PVINs receive, (2) a pruning of the synapses of PVINs from their postsynaptic targets, or (3) a change in the intrinsic electrical properties of PVINs. The latter is supported by recent evidence indicating that PVINs from nerve injured mice have a decreased input-output curve (Boyle et al., 2019; Qiu et al., 2022). In agreement with previous reports, we found a reduction in the intrinsic excitability of PVINs after nerve injury. But unlike in Boyle et al. 2019, we found an increased number of PVINs that fired adaptively and transiently following nerve injury. This discrepancy is possibly due to the different nerve injured models used (Corder et al., 2010; Boyle et al., 2019), but it remained to be established if transecting different sciatic nerve branches will result in the disparate changes in neural excitability. The deficiency in temporal and frequency encoding of PVINs is likely responsible for the observed decrease in their feed-forward synaptic efficacy on their postsynaptic targets (Cao et al., 2022). Such decreased PVINs output would ultimately undermine their ability to prevent the *Aβ* fiber-mediated activation of spinal pain pathways.

The inability of PVINs to maintain fast spiking was shown to accompanied by a lengthening in the time course of individual APs (slower AP rise time, half-width, fall time, and AHP time) and a clear potentiation of the medium component of the after-hyperpolarization potential, a phenotype mediated in part by SK channels (Sah and Faber, 2002). The sodium channel possibly contributes to the spike waveform changes along the spike train via its inactivation but is unlikely to explain the difference between naïve- and CCI- PVINs, as the spike thresholds were not affected following CCI (**Fig. 1G**). The involvement of SK channels, on the other hand, was confirmed by showing that their inhibition by the selective blocker apamin partially restored tonic firing in CCI-PVINs (**Fig. 6**). Our results suggested an increased contribution of SK current following nerve injury which leads to a reduction in the excitability of PVINs.

Our study does not exclude the possible contribution of ion channels, expressed in PVINs but not included in our models, in modulating the electrical activities of these neurons. The effect of hyperpolarization-activated cation current I_h_ were evident in both naïve- and CCI-PVIN recordings (**Table 4**). The voltage-gated subthreshold currents, including M-type potassium current I_M_ (Shah et al., 2008), the persistent sodium current I_NaP_ (Vervaeke et al., 2006), and the inward rectifier potassium current I_Kir_ (Akyuz et al., 2022), has been associated with generating mAHP and spike frequency adaptation. Despite the fact that these currents also modulate the passive membrane properties, which remain unaffected following nerve injury, their contribution to the PVINs excitability remains to be determined.

Our model is limited in assuming that PVINs are a homogenous population. This is not the case as only ∼60% of PVINs in lamina IIi and III are inhibitory (Qiu et al., 2022) and the inhibitory PVINs can be further clustered based on single nuclei RNAseq databases (Häring et al. 2018; Russ et al. 2021; Sathyamurthy et al. 2018). It is possible that each cluster of PVINs expresses its own complement of ion channels, which would be responsible for some of the variability that we observed in our experimental data when we put all our recordings from PVINs in one group (**Fig. 1 C-D, and M-N**). For instance, when compared to naïve-PVINs, CCI-PVINs consistently showed faster frequency adaptation (**Fig. 1D**), yet ∼20% neurons displayed persistent spiking during the 1- s long step current stimulation (**Fig. 1C**).

Our model showed that reducing cytosolic calcium-binding protein gradually elevates cytosolic calcium levels in PVINs (**Fig. 5B**) along the spike train. This is contrary to a previous model of PVINs (referred to as “Bischop model” hereafter) (Bischop et al., 2012). While the Bischop model and our model both showed that the low calcium buffer condition results in a higher amplitude of calcium transients, our model differed in the resulting calcium levels during the AP repolarization. The Bischop model described the role of cytosolic calcium buffering as a “calcium carrier”: during depolarization, the carrier binds to calcium ions and retains them temporarily; during the repolarization, cytosolic calcium levels decrease to their basal level rapidly (at a rate of 1 ms^-1^), causing the carrier to release calcium ions and “deliver” them to activate SK channels. Accordingly, increasing calcium buffer facilitates repolarization through the transient activation of SK channels, and thus increases the firing rate. However, based on the experimental data (Thayer and Miller, 1990; Muri and Knopfel, 1994), we decreased the calcium recovery rate *γ* to 0.001 ms^-1^ (See Material and Methods). Under this condition, the calcium buffer functions as a “chelator”: as the free calcium gradually accumulates along the spike train, the buffer effectively maintains cytosolic calcium at a lower level to prevent the activation of SK channels. The precise value of the calcium recovery rate is therefore important for the accuracy of the model and is required to be determined experimentally. Finally, our model shows that the endogenous calcium-binding proteins expressed in PVINs regulate neural excitability, and it will be of interest to examine the expression level of specific calcium buffers, such as parvalbumin, in naïve and nerve-injured conditions.

Our model suggests that the dysregulation of calcium dynamics is the primary cause for the abnormal activation of SK channels in CCI-PVINs. However, our proposed calcium-dependent mechanism is not the only way to regulate the activity of SK channels. For instance, our model showed that an increase in the calcium sensitivity of SK channels can also evoke spike frequency adaptation comparable to that of CCI-PVINs (**Fig. 3C-D**), which is regulated by the phosphorylation of the PIP_2_-CaM binding site of SK channels by the protein kinase CK2 (Bildl et al., 2004). Therefore, the modulation of SK channel activity by phosphorylation remains an important regulatory pathway that needs further investigation.

Our slow-fast analysis of the CCI-PVIN model established a significant role for intracellular calcium in controlling the firing behaviors of PVINs (**Fig. 7-8**). We showed that the slow dynamics of the intracellular calcium levels drive the state transition of the membrane potential in-between the stable steady state and the stable periodic oscillation state. The two-parameter bifurcation analysis further revealed how the patterns of such state transitions vary as different values of step currents are applied, which agrees well with the experimental observations from CCI-PVIN recordings. The two patterns of state transitions identified in our study: on-off switching and damped oscillations, have also been observed in different types of neurons in other areas of the central and peripheral nervous systems, where they serve unique physiological functions (Tal and Eliav, 1996; Wu et al., 2001; Xing et al., 2001; Amir et al., 2002; Clay and Shrier, 2002; Paydarfar et al., 2006; Århem et al., 2006; Huaguang et al., 2015; Zhao et al., 2020). Furthermore, our results on state transitions suggested that once the CCI-PVIN reaches the resting state within the 1-s depolarizing pulse, it can recover and resume spiking, which was verified in our preliminary prolonged CCI-PVINs recordings (data not shown). These findings indicate that the state transition of CCI-PVINs depends on the dynamics of intracellular calcium levels as well as the amplitude of the excitatory signals these neurons receive.

The PVIN model of reduced calcium buffer concentration consistently showed lower firing output when stimulated by either a stable step current or by a randomized synaptic current resembling that from Aβ fibers. These findings are relevant to pathological conditions that cause mechanical allodynia after nerve injury. When embedding our PVIN model into a simplified neural circuit model, the reduced firing output of PVIN failed to suppress the activity of the PKCγ HH type model, which can potentially activate the downstream pain-associated pathway. Despite the use of the simplified neural circuit model with a limited set of neurons and presynaptic inputs, our results demonstrate how changes in the intrinsic electrical properties of PVINs are involved in regulating the neural circuit dynamics and potentially, the high-level behaviors.

In conclusion, our study provided a calcium-dependent explanation for the decreased firing output of PVINs after nerve injury. It highlights the important roles of intracellular calcium dynamics in regulating the firing behavior of PVINs and identified SK channels as interesting therapeutic targets for treating the symptoms of mechanical allodynia. Given the heterogeneity of the PVINs, future work should develop an electrophysiological, and transcriptomic profiling of the PVINs in the superficial dorsal horn. Ion channels expressed in the different subsets of PVINs can also be added into our current model to better understand their diverse electrical activity.

## Acknowledgments

This work was supported by a project grant from the Canadian Institutes of Health Research to R.S.N. (CIHR PJT-162404) and the Natural Sciences and Engineering Council of Canada (https://www.nserc-crsng.gc.ca/index_eng.asp) discovery grant to A.K. and the McGill Initiative in Computational Medicine (https://www.mcgill.ca/micm/) team grant to A.K. and R.S.N. X.M. was supported by the Globalink Graduate Fellowship. The authors would like to acknowledge the support of Dr. Hugues Petitjean and Ms. Francesca Leonardo for their contribution to an earlier set of experiments not included in this manuscript.

## Author contributions

X.M., L.M., R.S.N., and A.K. designed the research project; L.M. and H.Q. designed and performed the data collection of electrophysiology experiments; X.M. and A.K. designed the model and analyzed it; X.M. analyzed data and performed the computation; X.M wrote the first draft of the paper; X.M., L.M., R.S.N., and A.K. edited the paper; X.M. wrote the paper; R.S.N. and A.K. conceived the original idea, supervised the project and obtained the funding; X.M. performed the research.

---

**Extended Figure 2-1.**
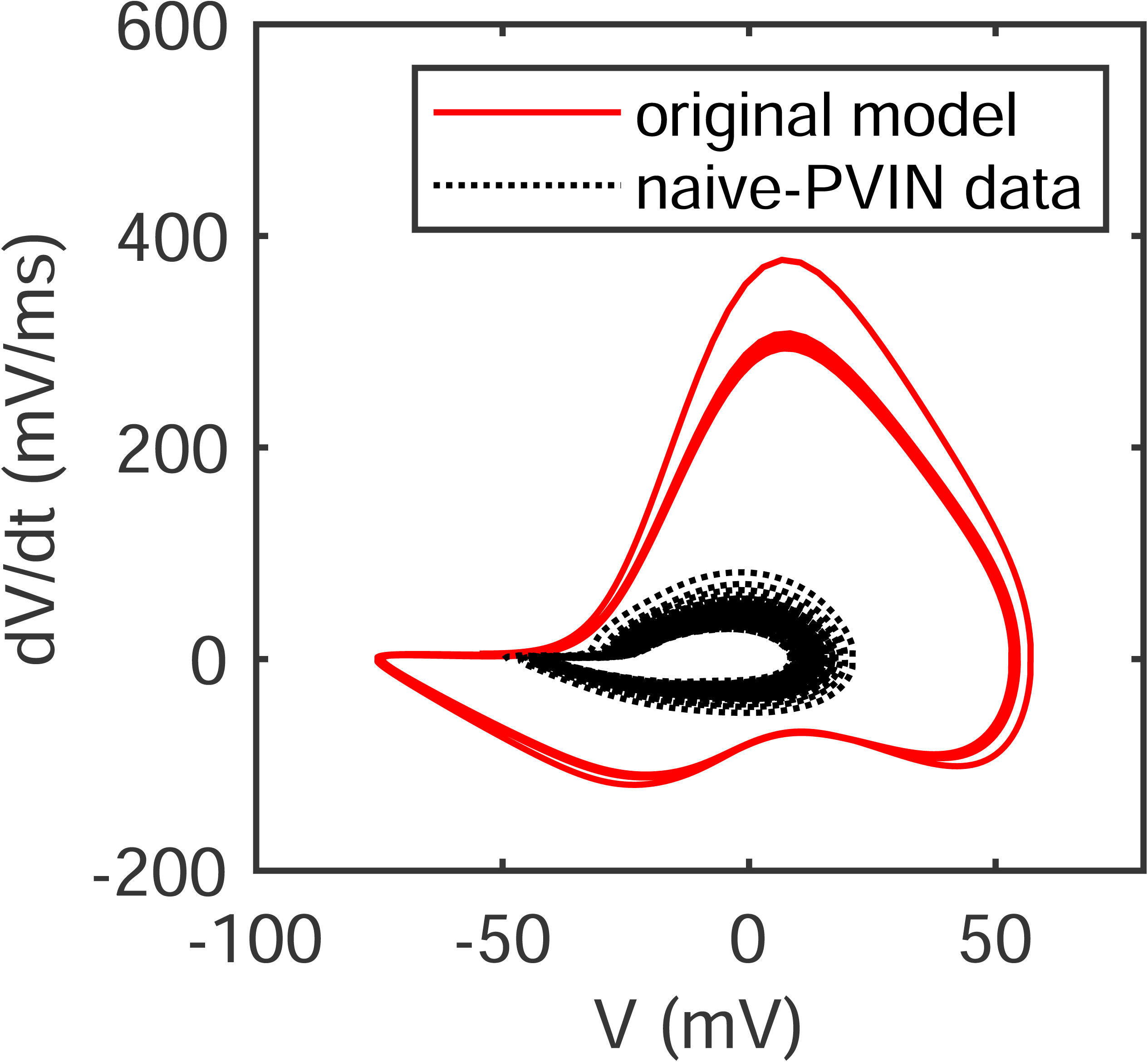
The AP cycles of naïve-PVIN recordings differ significantly from those generated by the original model developed in (Bischop et al., 2012). The AP-cycles generated by the model (red solid curve), with its original parameters, have very few common features with the naïve-PVIN recording (black dotted curve) when plotted in the (V, dV/dt)-plane. (The voltages are not corrected for liquid junction potential in this figure).

**Extended Figure 2-2.**
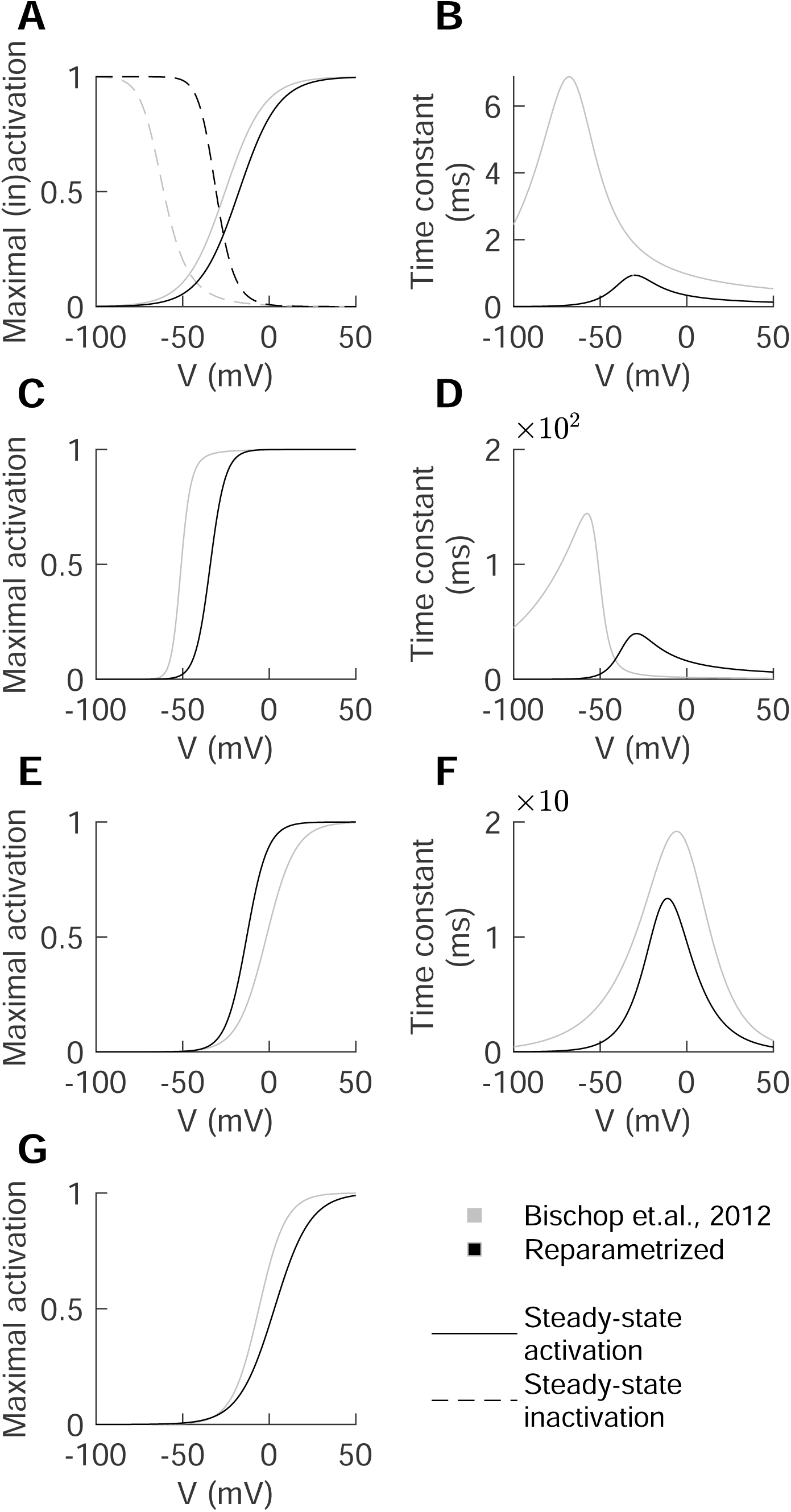
The kinetics of ion channels before (grey) and after (black) reparametrization of the Hodgkin-Huxley model of PVIN when fitted to electrophysiological data. **A-B**, The steady-state activation (solid line), inactivation (dashed line) and time constant of sodium current; **C-D**, The steady-state activation and time constant of Kv1 current. **E-F**, The steady-state activation and time constant of Kv3 current. **G**, The steady-state activation of Ca current.

